# A new iterative framework for simulation-based population genetic inference with improved coverage properties of confidence intervals

**DOI:** 10.1101/2024.09.30.615940

**Authors:** François Rousset, Raphaël Leblois, Arnaud Estoup, Jean-Michel Marin

## Abstract

Simulation-based methods such as approximate Bayesian computation (ABC) are widely used to infer the evolutionary history of populations from molecular genetic data. We describe and evaluate a new iterative method of statistical inference about model parameters, which revisits the idea of inferring a likelihood surface using simulation when the likelihood function cannot be evaluated. It is based on combining the random forest machine learning method, and multivariate Gaussian mixture (MGM) models, in an effective inference workflow, here used to fit models with up to 15 variable parameters. In addition to the traditional assessment of precision in terms of bias and mean square error, we also evaluate the coverage of confidence intervals. The method is compared with approximate Bayesian computation using random forests (ABC-RF), a non-iterative method sharing some technical features with the proposed approach, across scenarios of historical demographic inference from population genetic data. It is also compared to another iterative method, sequential neural likelihood estimation (SNLE). These comparisons highlight the importance of an iterative workflow for exploring the parameter space efficiently. For equivalent simulation effort of the data-generating process, the new summary-likelihood method provides intervals whose coverage is better controlled than the marginal coverage of intervals provided by ABC with random forests, and than generally reported for ABC methods. The iterative workflow can also yield greater improvements in estimator precision when larger datasets are used.

## Introduction

The likelihood function is a classical component of efficient statistical inference methodologies. In instances where the likelihood function cannot be computed in a reasonable time frame, alternative statistical methods can be employed. These include moment-matching techniques when the analytical results for moments of a response variable are known, or approaches based on the simulation of the putative data-generating process. Among the latter simulation-based methods, approximate Bayesian computation (ABC) has been particularly developed, notably for applications in population genetics (Beaumont, 2010; Beaumont *et al*., 2002; Bertorelle *et al*., 2010; Tavaré *et al*., 1997), but also in diverse other fields of biology (Schälte and Hasenauer, 2020) and beyond (e.g., Akeret *et al*., 2015; Sisson *et al*., 2019). In molecular evolutionary studies, ABC has been widely used to infer the past history of migration, founding events, invasion routes and introgressions among populations (e.g., Fraimout *et al*., 2017), but also selection pressures (e.g., Nakagome *et al*., 2015) or genomic rearrangement rates (e.g., Moshe *et al*., 2022).

The idea of estimating the likelihood function by simulation is less developed, although it goes back at least to Diggle and Gratton (1984). Some recent machine-learning methods, such as sequential neural likelihood estimation (SNLE, Papa-makarios *et al*., 2019), incorporate this step, although they perform inference via the posterior distribution rather than the likelihood function. Diggle and Gratton’s more strictly likelihood-based approach has had limited follow-up in the form of practically implemented likelihood methods for more complex models, particularly when the data are represented by many descriptive statistics (i.e., “summary statistics”). The “synthetic-likelihood” method (Wood, 2010) may be seen as derived from Diggle and Gratton (1984), except that it assumes the summary statistics to have a multivariate normal distribution for each parameter value. An extension of Diggle and Gratton’s approach not making this assumption has been described by Rousset *et al*. (2017). As this is a likelihood inference using the information retained in the summary statistics, we refer to this method as “summary-likelihood (SL) inference”. Summary likelihood is not full-data likelihood but is still a form of likelihood, which one can evaluate if the full data have been thrown away and only some summary statistics have been retained. When the number of summary statistics is large, inferring the likelihood surface becomes more tractable by applying dimension-reduction techniques. Various machine learning methods can be employed for this purpose; here, we adopt the random forest approach (Breiman, 2001; Geurts *et al*., 2006). This method is also used in ABC-RF (Collin *et al*., 2021; Pudlo *et al*., 2016; Raynal *et al*., 2019).

The aforementioned methods all address the issue of inferring the parameters ***θ*** of a hypothetical data-generating process, utilizing simulations of this process for varying values of the vector ***θ***. In Rousset *et al*. (2017) as in Diggle and Gratton (1984) and Wood (2010), values of ***θ*** are drawn and for each draw, the distribution of summary statistics is estimated using a moderately large number of simulations of the data-generating process. In this study, we present a more efficient summary-likelihood inference workflow that requires significantly fewer simulations to provide accurate inferences. This iterative workflow facilitates the inference of more parameter-rich models, even when simulating observations is computationally expensive. The new workflow employs a single simulated observation for each drawn value of ***θ***. This approach aligns with the general methodology of ABC methods, where each drawn ***θ*** and the corresponding summary statistics for each simulated observation form one line of a table commonly known as the reference table. The new iterative method has already been used to fit a simple two-parameter model of evolution of experimental populations by Laugier *et al*. (2025).

The idea of estimating the likelihood function for summary statistics from an ABC reference table already appears in the ABC literature (e.g., Fan *et al*., 2013; Papamakarios *et al*., 2019; Rubio and Johansen, 2013), but this has not yet led to accessible and validated software implementations of summary-likelihood inference. Indeed, a potentially substantial drawback of this idea is that inferring the likelihood surface in high dimensions is a complex task. For inference of likelihoods or joint full posterior distributions, the importance of iterative (or “sequential”) methods of construction of the reference table has been repeatedly stressed (e.g. Blum and François, 2010; Cranmer *et al*., 2019; Del Moral *et al*., 2006; Lueckmann *et al*., 2021; Toni *et al*., 2009). Modern variants employ neural networks for density estimation (Blum and François, 2010; Greenberg *et al*., 2019; Lueckmann *et al*., 2017; Papamakarios and Murray, 2016; Papamakarios *et al*., 2019). However, in practice, non-iterative ABC methods (e.g., Beaumont *et al*., 2002; Blum and François, 2010; Pudlo *et al*., 2016) remain widely used. Such methods can, in principle, enable amortized inference, whereby a single reference table (or even a single trained neural network) can be used to analyze many datasets. This can yield substantial computational savings when data are repeatedly generated under the same design (e.g., Zammit-Mangion *et al*., 2025). However, in many real-world applications of the non-iterative ABC methods described above, datasets differ enough in design or characteristics that such amortization is not feasible.

ABC methods are employed for the purpose of inferring posterior distributions for parameters, given a prior distribution for ***θ***. Subsequently, credible intervals can be derived from the posterior distribution. As an alternative approach, likelihood-ratio based confidence intervals can be computed when a likelihood surface is inferred. The intervals returned by the different methods can be compared in terms of coverage, which is the probability that the interval contains a data-generating parameter value. It is anticipated that credibility intervals will provide correct coverage on average across the prior distribution of the parameters, while confidence intervals are constructed to ensure correct coverage for any possible value of the parameters: for different perspectives on these two concepts (credibility and confidence intervals) see for example Neyman (1977) or Casella and Berger (2002). However, it appears that a significant number of ABC methods are unable to effectively control the prior-averaged coverage of credibility intervals. For instance, Raynal *et al*. (2019) evaluated coverage of credibility intervals for ABC-RF as well as for basic rejection ABC and its elaborations using adjusted local regression (Beaumont *et al*., 2002), ridge regression (Blum *et al*., 2013), or neural networks (Blum and François, 2010). They found that ABC-RF intervals were conservative, with 100% coverage for 95% nominal probability. Additionally, they found that the coverage of other ABC methods was contingent upon a required rejection threshold, which is challenging to accurately set a priori. Consequently, they also generated anti-conservative intervals for certain threshold values. As observed by Hermans *et al*. (2022), the issue of coverage control has been seldom addressed in earlier machine learning literature. The deep learning-based methods they examined were all found to produce anti-conservative intervals, indicating a clear need for improvement in the coverage of intervals in simulation-based inference methods. Consistent with this, coverage of credibility intervals has received increasing attention in recent work (e.g., Frazier *et al*., 2024), with some studies explicitly distinguishing the coverage properties of Bayesian credible intervals from those of frequentist confidence intervals (Dalmasso *et al*., 2024).

In this paper, we present an evaluation of the performance of our new summary-likelihood workflow using simulation scenarios of inference of demographic history by analysis of population genetic data. A toy simulation scenario completes the tests of the method. Compared with the non-iterative ABC-RF method, our results underscore the value of an iterative workflow for improving inference when accurate exploration of the parameter space is critical. We show that the new workflow allows more uniform control of the coverage of intervals than previously reported for ABC methods. The main deviations from ideal control appear due to lack of information about parameters in the data, a common feature of demo-genetic inferences, and of other fields of application of simulation-based inference (e.g., Auger-Méthé *et al*., 2021; Daly *et al*., 2018; Fan *et al*., 2019). We also provide some comparison with the SNLE workflow implemented in the `sbi` package (Tejero-Cantero *et al*., 2020). This iterative workflow uses highly efficient neural network methods to infer likelihood surfaces and posterior densities, and provides credibility intervals. For problems of higher dimension than those considered here (up to 15 parameters), this method appears faster. However, our simulations show that, in the population-genetic scenarios examined here, the coverage of the intervals provided by SNLE is not always well controlled.

## Material and Methods

### The inference workflow

Starting from a reference table built from a limited number of simulations, an initial estimation of the summary-likelihood surface is derived. Then, new parameter points are sampled with greater probability in regions of high inferred likelihood, with the objective of more accurately inferring the likelihood surface in such regions. These sampling and likelihood-surface estimation steps are repeated iteratively to augment the reference table and to obtain progressively more precise inferences of the likelihood surface.

In this Section we provide a first description of these steps, introducing terminology and notation. For clarity, the simulation results included in the reference table are called “samples” to distinguish them from the “data” to be analyzed, whether the latter are real data or simulated ones. In the same way, we distinguish the data-generating parameters values (denoted ***θ***^*†*^), which are not information used by the inference workflow, from the sample-generating parameters, which are essential information included in the reference table. Following well-established notation that we will repeatedly use below, probability densities will be denoted as P, with indices denoting the random variables whose values are the arguments of the function. In particular, P_*Y* ;**Θ**_ denotes the density of *Y* values (where these values may represent the data, or some simulated sample) as function of parameter values **Θ**, and P_*Y*,**Θ**_ denotes the joint density of *Y* and **Θ**, under the assumption that **Θ** is sampled from some distribution. P_*Y* ;**Θ**_(*S*; ***θ***) is thus the probability (or probability density) of a sample *S* for a given parameter vector ***θ***.

#### Inferring the likelihood from a reference table

The likelihood *L*(***θ***; 𝒟) of ***θ*** given observed data 𝒟 is generally defined, up to a constant factor, as P_*Y* ;**Θ**_(𝒟; ***θ***), viewed as a function of the parameters for fixed data. Given a distribution *i*_**Θ**_(***θ***) for the parameters, the joint distribution of samples *S* and parameters ***θ*** can be written P_*Y*,**Θ**_(*S*, ***θ***) = *i*_**Θ**_(***θ***)P_*Y* ;**Θ**_(*S*; ***θ***), and the likelihood can be written as

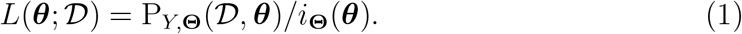

Accordingly, to estimate the likelihood function from the reference table, one can first estimate a joint density P_*Y*,**Θ**_ of samples and parameters, from their realized joint distribution in the reference table. One can then divide the value in *Y* = 𝒟 of this estimated joint density function by an estimate of the marginal parameter density function *i*_**Θ**_, deduced from the joint density, to obtain the likelihood function. Since the term “instrumental distribution” is commonly used to refer to a probability distribution used in an algorithm for the estimation of a target quantity, such as a likelihood or posterior distribution, we thus view the **Θ** vectors in the reference table as samples from an instrumental distribution which is estimated conjointly with the distribution of summary statistics in the reference table.

As in Papamakarios *et al*. (2019), the iterative proposal mechanism for new parameter points does not asymptotically bias likelihood learning. In the present workflow, new parameters points are sampled in each iteration in order to preferentially fill the region of parameter space with a high likelihood (as detailed below, Section Refinements of likelihood surface inference through iterations). The joint and marginal densities, and the likelihood, are re-estimated in each iteration. After a few iterations, the inferred *i*_**Θ**_(***θ***) is usually quite different from the distribution of parameters used to construct the initial reference table.

#### From raw statistics to projected statistics

Here, as in most applications of ABC, the information provided to the inference method is typically a vector of statistics **u**(𝒟), summarizing higher-dimensional observed data 𝒟 which is the information available to the analyst. For example, the largest genetic datasets considered in our simulation study involve genetic information at 10,000 genetic markers from 120 individuals, which is summarized by 130 statistics; but in some applications **u**(𝒟) may just be a vectorial representation of the full 𝒟. The simulated samples for each drawn ***θ*** must then also be expressed as a vector of summary statistics **u**(*S*). As with ABC, it is up to the user to provide statistics that are informative for a given inference problem.

In our SL method, the potentially large number of *raw* summary statistics describing the data or the simulated samples in the reference table (up to 130 summary statistics in our simulations), is typically reduced by a *projection step* to a smaller number of *projected* summary statistics **p**[**u**(*D*)], in order to reduce the dimension of the joint distribution, of parameters and of statistics, which is estimated. “Projection” here refers to the idea of non-linear projection to a lower-dimensional space. We obtain the projected summary statistics by performing a non-parametric regression of each parameter *θ*_*j*_ on the raw summary statistics. This regression is performed using a variant of the random-forest method, combining features of its original version (Breiman, 2001) and of “extremely-randomized trees” (Geurts *et al*., 2006). Various other projection methods could be used in SL, provided they provide predictions that avoid overfitting. The random-forest method was retained for the same reasons as in ABC-RF: it is fast, efficient, does not require ad-hoc adjustment of control settings for each application, and the out-of-bag predictions can be used to easily avoid overfitting (e.g., Breiman, 2001, p.11; Hastie *et al*., 2009). It thus fulfills the need for a method which can be applied automatically in any inference problem (Raynal *et al*., 2019). This choice also facilitates comparison with ABC-RF, as differences between the performance of ABC-RF and summary likelihood cannot be attributed to the choice of widely different projection methods.

In our use of random forests, a joint vector **p**[**u**(*S*)] of projected summary statistics is defined, for each simulated sample *S*, as the vector of predictions 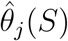 of each element *θ*_*j*_ of the ***θ*** parameter vector by its random-forest regression. This reduces the number of statistics to the number of parameters, which is the minimum required for identifiability of the parameters. Additional statistics can be retained insofar as computations do not become impractically slow, but have not been considered here. Fearnhead and Prangle (2012) already advocated summarising a sample by the posterior expectations of each parameter given that sample, which can be approximated in practice using random-forest regression. Here, however, the random-forest regression targets posterior expectations under the instrumental distribution induced by the iterative algorithm, rather than under any fixed, pre-specified prior.

An initial reference table is thus constructed as follows. Parameter vectors are drawn from the initial instrumental distribution 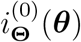 for parameters, and one sample *S* (i.e., one realization of the biological and sampling process) is drawn for each parameter vector. Raw summary statistics **u**(*S*) are computed, and projected statistics **p**[**u**(*S*)] are deduced from them for each such sample.

#### Estimating the summary-likelihood function

The likelihood function for projected statistics is written

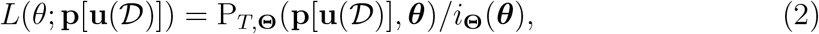

in terms of the joint density P_*T*,**Θ**_ of projected statistics and parameters.

To model this joint density from the reference table, we use by default a Multivariate Gaussian Mixture (MGM) model. MGMs have previously been used to infer posterior distributions of parameters (Bonassi *et al*., 2011), and a related approach, Gaussian Locally Linear Mapping (GLLiM, Deleforge *et al*., 2014), has also been used in simulation-based inference (Häggström *et al*., 2024). We also considered Masked Autoregressive Flows (MAFs, Papamakarios *et al*., 2017), a deep-learning approach for estimating unconditional or conditional probability densities. In our first attempts to use MAFs, they were trained directly on the final reference table. Because this is substantially slower than fitting MGMs, MAFs did not appear suitable as a systematic replacement for MGMs in our workflow. In practice, MAFs are more efficiently trained sequentially, updating the inferred density at each iteration by warm-starting from the previous iteration; this is the strategy used in SNLE, whose performance will be compared to summary-likelihood for a set of selected simulation scenarios.

Once a joint density estimate 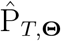 is obtained using MGM models, the likelihood for any ***θ*** is estimated by

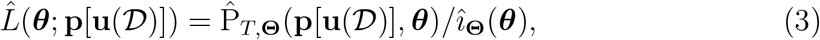

where, when the joint density estimation uses MGM models, the estimate *î*(***θ***) of the instrumental density is easily deduced from the joint density estimate by marginalization of the latter density over the projected statistics. One may think that such estimation is not needed, at least in the first iteration when the instrumental distribution typically has some simple known form. However, in subsequent iterations such knowledge is not available because the instrumental density (as automatically generated by rules discussed later) is implicit and complex.

When Masked Autoregressive Flows are used, the likelihood function can be directly estimated by training a conditional MAF (Papamakarios *et al*., 2017, Section 3.4; Papamakarios *et al*., 2019, in order to learn the conditional density of samples given any ***θ*** values, on the joint distribution of parameters and statistics in the reference table.

Maximum summary-likelihood estimates (summary-MLEs or MSLEs, denoted 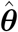) are deduced from the inferred likelihood surface, by numerical maximization with respect to ***θ*** of the estimated likelihood as given by eq. 3. Summary-likelihood ratio tests (summary-LRTs) are also deduced from the likelihood surface. In particular, when testing a value *θ*_*i*_ of a given element *i* of ***θ***, we compute the constrained maximum of the estimated summary-likelihood surface given *θ*_*i*_ (yielding the constrained maximum summary-likelihood estimates 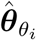), and compare it to the global summary-likelihood maximum. Thus, we use the profile likelihood (e.g., Davison, 2003, Section 4.5.2) for all likelihood-ratio tests, with test statistic

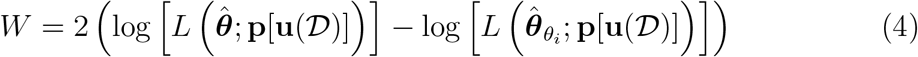

and p-value given by the tail probability P(*X > W*) for a *χ*^2^-distributed variable with number of degrees of freedom equal to the number of fixed parameters in 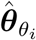, i.e. 1 in the present applications of the test.

#### Refinements of likelihood surface inference through iterations

In each iteration, new parameter values are drawn, simulations of the process are performed and added to the reference table, and the above steps of the workflow are repeated on the incremented reference table. The sampling of parameters should still permit further exploration to prevent entrapment in the initial high-likelihood region, which may be disjoint from the final one (as will be illustrated by the results on the 7-parameter human admixture scenario). The implemented sampling procedure further allows exploration of the parameter space beyond the previously sampled ranges, and users can specify “absolute” bounds that should not be exceeded during such exploration. The software also allows users to specify not only ranges for each parameter, but also arbitrary additional constraints on any combination of parameters (this is used in some of our simulations).

The rules for drawing *n*_*i*+1_ new parameter vectors after iteration *i* are detailed in the Supplementary Information (Section S.1.1). They are defined to preferentially fill the region of parameters with a given minimum likelihood ratio *l*_min_ relative to the current summary-MLEs. In practice, *l*_min_ was set to the 95% threshold for two-dimensional confidence regions. The sampling also allows exploration beyond the boundaries of this “top” region. The sampling step within the “top” aims to fill it uniformly rather than in proportion to the likelihood ratio.

The iterative workflow allows the projections to be re-computed. However, random-forest computations on large tables take time, so some elaborations have been implemented to shorten them. First, for large reference tables, they use a subset of the reference table, mostly defined from points identified as belonging to the top of the likelihood surface (see Supplementary Material for details). Second, projections are not re-computed when more than 90% of the selected points were already used in the previous computation of the projections. The easiest way to accelerate inferences in parameter-rich models without substantially compromising performance may be to reduce this threshold so that projections are re-computed less often.

#### Implementation

All the above-described steps of summary-likelihood inference have been implemented in an automated workflow in the `Infusion` R package, which calls the `ranger` R package (Wright and Ziegler, 2017) for random-forest methods, and the `Rmixmod` R package (Lebret *et al*., 2015) for MGM modeling. .

### Control of inference workflow

Our workflow potentially depends on many control parameters, but we set default values for all of them, which were used for all simulations in this paper, unless mentioned otherwise. These values, as described here and further detailed in Supplementary Section S.1, are automatically selected by our implemented procedures unless users specifically request different values.

In particular, for fitting *n*_p_ parameters to a given dataset, a final number of 1000(3*n*_p_ − 1) sample simulations are run. Supplementary Table S.1 illustrates computations times for single datasets for different simulation designs, and Supplementary Section S.1.6 provides further details on how the number of samples added at each iteration is controlled. Smaller references tables appear to lead to degraded performance (one example being given in Supplementary Table S.15) and are therefore not recommended. Larger tables may require substantial increases in computation times, which may be a small concern when only one dataset is analyzed, but would be unpractical when inferences are performed on series of hundreds of simulated datasets.

While the final size of the reference tables was predefined in our simulation study, alternative and more adaptive stopping rules could be used, based in particular on two criteria implemented in the R package. The first criterion is the estimated precision of the log profile likelihood ratio statistics at the bounds of the inferred confidence intervals. A bootstrap procedure is implemented to evaluate root mean square errors (RMSEs) of prediction of these log-ratio statistics. It is thus possible to request termination of iterations when these RMSEs, averaged over the different interval bounds, or when all such RMSEs, are below a certain threshold. A second criterion is the comparison of the distribution of samples simulated in two ways: samples from the simulator of the process being inferred, versus samples from the inferred distribution. These distributions can be compared as in a classifier two-sample test (Lopez-Paz and Oquab, 2017), by training a classifier to assign samples to either distribution: the better the inferred distribution, the lower the classification performance. This has previously been used to compare different methods of inference of posterior distributions (Lueckmann *et al*., 2021), but can be used here to compare the distributions of samples given specific values of the inferred parameters, such as their MSLEs.

#### SNLE inference

To perform SNLE inference, we used the Python package `sbi` version 0.25.0, with default controls for training the MAF neural density estimator. In particular, the flow consisted of five autoregressive transformations, each parameterized by a Masked Autoencoder for Distribution Estimation (MADE)-type neural network (Germain *et al*., 2015) with hidden layers of size 50 and two blocks. Dropout and batch normalization were disabled. The autoregressive networks used ReLU activation functions. Between successive transforms, random permutations were applied to increase flexibility of the flow.

Posterior samples were obtained using the default Markov chain Monte Carlo (MCMC) sampler implemented in `sbi`. This corresponds to slice sampling with multiple parallel chains. The default configuration uses 20 chains, with 100 warmup steps per chain and thinning factor equal to one.

Final reference tables had the same size as in our default summary-inference workflow, and we ran ten rounds of the sequential procedure with an equal number of simulated samples per round (Lueckmann *et al*., 2021, Appendix Section A.4).

### Design of simulation study

Our simulation study is mainly based on two scenarios of demographic history of populations which have been considered in previous developments of ABC with random forests. However, we also consider 15-parameter toy examples where the number of raw summary statistics is the number of parameters, so that projections are not needed and inferences are relatively fast.

#### Toy examples

In the toy examples, we estimate the covariance parameters of a 5-dimensional normal distribution. The data-generating values, and the parametrization of this model are detailed in Supplementary Section S.3.1. The summary statistics are defined as the 15 distinct elements of the observed covariance matrix of 50 draws from a multivariate normal distribution of dimension 5 with the given covariance matrix.

#### Origin of invasive ladybird populations

This scenario (Figure 1, left) was already taken as an example for ABC-RF analyses by Pudlo *et al*. (2016). It is motivated by the invasion of the ladybird beetle *Harmonia axyridis* in Europe, and by a dataset of genotypes at 18 microsatellite loci, from a total of 126 individuals from four populations (Lombaert *et al*., 2011). The fitted model has 8 parameters: effective population sizes *N*_1_ to *N*_4_, admixture time *t*_1_, admixture proportion *r*_a_, and two mutation parameters 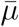 and 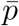. We estimated the composite parameters 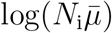 instead of the *N*_*i*_s. Further details on model, data, and sampled parameter ranges are given in Supplementary Section S.4.1.

**Figure 1:**
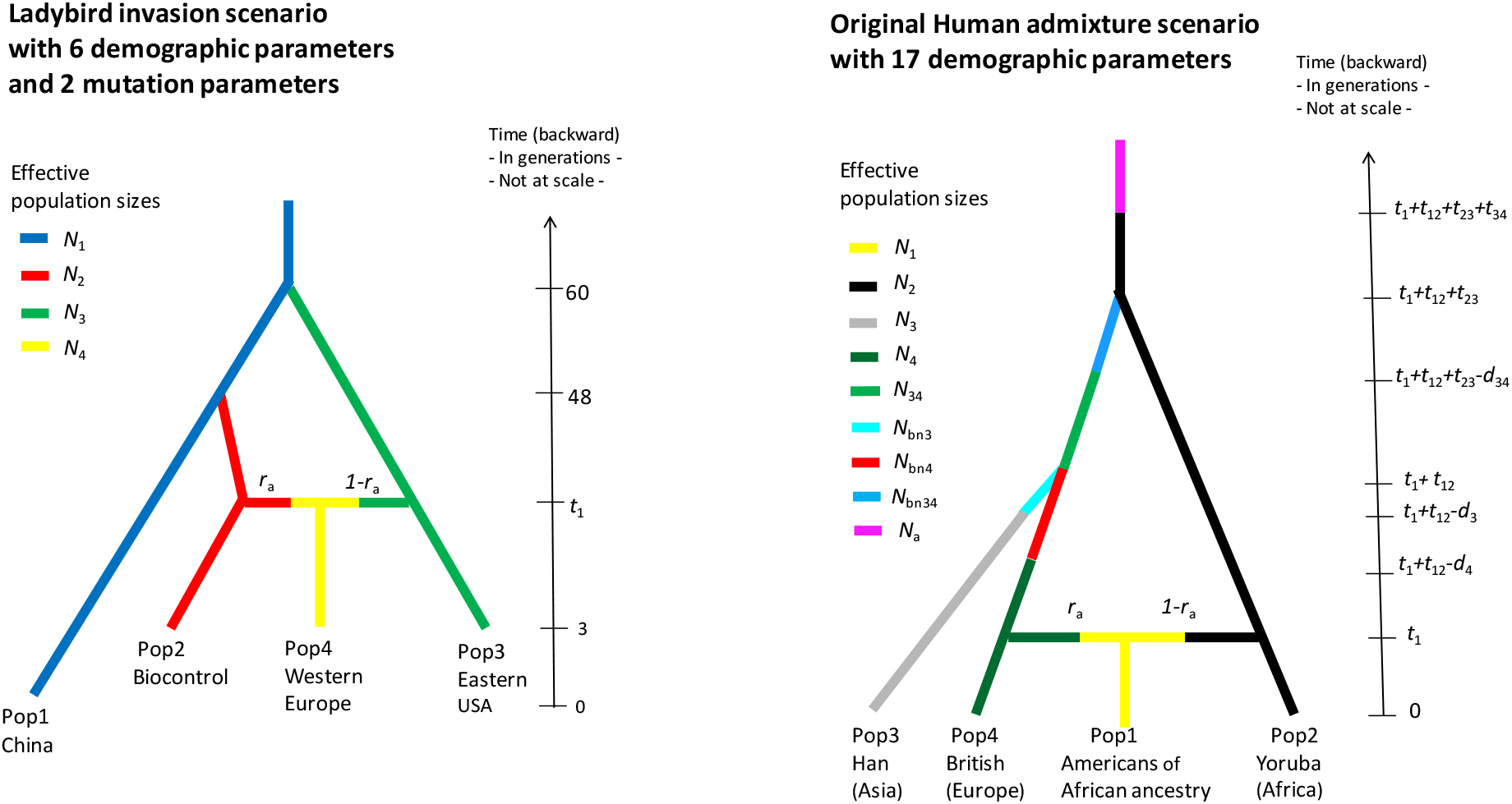
The two scenarios of historical demography. Left: ladybird invasion scenario; right: Human admixture scenario. See Text for description of parameters.

#### Human admixture scenario

For this second scenario (Figure 1, right), the model and data were also already considered in previous papers on ABC-RF (Collin *et al*., 2021; Raynal *et al*., 2019). It is a scenario of admixture between populations of European and African ancestry in America. More precisely, after an ancient demographic change at time *t*_4_ in the ancestral African population, an out-of-Africa colonization event occurs at time *t*_3_ that gives an ancestral out-of-Africa population which secondarily splits into one European population and one East Asian population at time *t*_2_. Finally, a genetic admixture event occurs between populations of European and African ancestry in America at the time *t*_1_, with a proportion *r*_a_ of European ancestry. Additional parameters *N*_bn3_, *N*_bn4_, *N*_bn34_ describe the population sizes during bottleneck events associated to colonization events along the population tree, and *d*_3_, *d*_4_, and *d*_34_ describe the duration of these bottlenecks. As detailed and justified in Supplementary Section S.5.1.1, we re-parametrized the model in terms of *t*_12_ = *t*_2_ − *t*_1_, *t*_23_ = *t*_3_ − *t*_2_, *t*_34_ = *t*_4_ − *t*_3_.

We explored different versions of this model, with 7 or 13 estimated parameters, by assuming that the values of other parameters were fixed and known. A real dataset of 5000 SNP markers (extended to 10000 in some simulations), genotyped in four human populations (The 1000 Genomes Project Consortium, 2012), defines the sampling design of the simulated data. The four populations include Yoruba (Africa), Han (East Asia), British (Europe) and American individuals of African ancestry.

Few parameters may be practically identifiable in the Human admixture scenario. For this reason, Raynal *et al*. (2019) and Collin *et al*. (2021) reported estimation performance for only a few parameters, with good performance only for the admixture rate *r*_a_ and for some composite parameters defined as ratios of population size to time interval parameters. However, such composite parameters complicate the interpretation of the results. For example, the parameter space is no longer defined only by box constraints (i.e., by the range of each parameter), and marginal prior distributions are no longer uniform. This implies that low relative RMSEs, which are defined relative to the marginal range of each parameter, are not necessarily indicative that the parameter is practically identifiable.

For these reasons, we considered composite parameters only in one of two considered 13-parameter variants of the Human admixture scenario. More specifically, in this variant we considered the following composite parameters, sometimes referred to as “bottleneck intensity” parameters in the colonization/invasion literature: *b*_3_ = *d*_3_*/N*_bn3_, *b*_4_ = *d*_4_*/N*_bn4_, and *b*_34_ = *d*_34_*/N*_bn34_. Estimating additional population size parameters specific to the branches of the tree that are affected by bottlenecks is difficult. In this simulation variant, we therefore do not try to estimate two such parameters, *N*_3_ and *N*_4_. Supplementary Table S.19 presents results for another 13-parameter variant without any composite parameters and with estimated *N*_3_ and *N*_4_.

Further details on model, data, and sampled parameter ranges are given in Supplementary Section S.5.1.1.

#### Assessment of confidence and credible intervals

The concept of confidence interval is based on the control of coverage whatever the data-generating parameter values ***θ***^*†*^, rather than only on average over a prior distribution, so we assess coverage for given ***θ***^*†*^ values, with only a few exceptions.

For fixed prior distributions, credibility intervals may become asymptotically equivalent to confidence intervals for “large samples”, i.e., when the information contained in the parameters increases indefinitely with sample size (Lehmann and Casella, 1998, Section 6.8). Credibility intervals and confidence intervals may then be seen as asymptotic approximations to each other. But this asymptotic equivalence may fail when (i) coverage is evaluated conditionally for a parameter at the boundary of the prior distribution and (ii) for some definition of posterior intervals. In particular, intervals based on central quantiles of the posterior distribution (“central posterior intervals”) will never cover the ***θ***^*†*^ value when it is at a boundary.

This boundary effect should not be overlooked: (i) it may be widely ignored by practitioners (reports of performance focusing on good marginal coverage encouraging such ignorance), and (ii) it is a real concern in practice insofar as software (including ABC-RF) often report central posterior intervals, which leads to the exclusion of parameter values at range bounds from reported intervals, even when there is actually not enough information in the data to actually reject such values. For the following comparisons, we implemented the computation of highest posterior density (HPD) intervals from the ABC-RF output, which are expected to have better coverage at the boundaries. For unimodal posterior distributions, these are also the shortest credible intervals.

In principle, assessment of performance of confidence intervals calls for estimation of coverage for many different ***θ***^*†*^ values. However, this would be unpractical here due to the high-dimensional parameter space and the computational cost of the simulations. Supplementary Section S.2 details the rules used to select several data-generating parameter values ***θ***^*†*^; and Supplementary Table S.5 and Section S.5.1.2 show the values used for the ladybird invasion scenario and the Human admixture scenario, respectively.

### Performance summaries

We evaluated the bias and root-mean-square error (RMSE) of the following point estimators: the summary-MLEs, and the mean and median of the posterior distributions for ABC-RF. Each Figure and Table for given data-generating parameter values is based on running the inference workflow on 400 simulated datasets (but 200 or 1000 datasets for some of the Supplementary Tables and Figures).

Parameter transformations were applied, mainly in order to homogenize the RMSEs of the estimators of transformed parameters in the population genetic scenarios. In practice this means that population size and mutation rate parameters were log-transformed, but for simplicity we also applied the following automatic transformation rule to other parameters based on their explored ranges: if upper bound is ≥ 500, log(1+.) or log(.) transformation is used, depending on whether the lower bound is zero or not. All logarithms are base 10 logarithms. ABC-RF inferences used uniform priors on the transformed scales.

The RMSE summaries may not lead to clear conclusions. Which of ML and posterior estimates have lower RMSE depends on the location of data-generating values ***θ***^*†*^ relative to the prior distribution used in a Bayesian inference (e.g., Casella and Berger, 2002, p. 333), so we do not expect RMSE criteria to systematically favor one class of estimators over the other when examined conditionally given ***θ***^*†*^ values. Moreover, an efficient posterior estimate is expected to have lower RMSE on average when ***θ***^*†*^ values are chosen randomly in their prior distributions. Yet, systematic departures from such theoretical expectations can occur if one of the methods does not locate well its point estimates, as a result of poorly inferring the likelihood function or the posterior distribution.

Bias and RMSE of estimators will be reported on a scale relative to the width of the explored ranges: for the *i*th element *θ*_*i*_ of the parameter vector, we use transformed parameter values *ϑ*_*i*_ as described above, and evaluate the following scaled means:

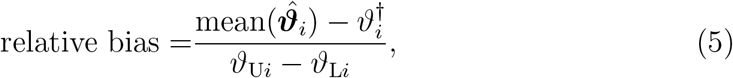

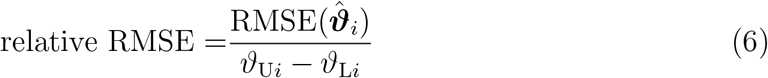

where 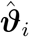 is the vector of estimates of *ϑ*_*i*_ over the simulated datasets; and 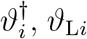, *ϑ*_L*i*_ and *ϑ*_U*i*_ are the corresponding data-generating value, lower, and upper bound of explored range of the transformed *i*th parameter, respectively. With relative variance defined as 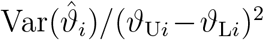, the standard decomposition of mean-square error as variance plus squared bias holds for these relative values.

This is useful in particular to compare the RMSE of a summary-likelihood estimator to that of a non-identifiable parameter whose estimator would be uniformly distributed, and would then have a relative RMSE equal to 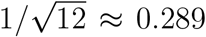 if the data-generating value were the mid-range of the parameter (and higher otherwise). Another useful diagnostic pattern for poorly identifiable parameters is the relative variance of summary-MLE versus posterior estimates: the posterior mean estimator should have low variance, as it should approach the prior mean for all datasets. Thus, although an efficient posterior estimate is expected to have slightly lower RMSE on average over a prior distribution, a markedly lower ratio of the variance of posterior estimates to the variance of summary-MLEs instead suggests that the parameter is not well estimated by ABC-RF.

The actual conditional coverage of nominal 95% intervals will be reported for both the confidence and credibility intervals. For each element *θ*_*i*_ of the parameter vector, a more informative summary, not depending on a conventional level such as 95%, is the actual distribution of p-values of the test whose null hypothesis is that 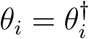. This distribution should be uniform in ideal conditions, and will be reported in Figures. We used the profile likelihood function for such tests, as previously described.

We also evaluate the performance of intervals provided by parametric bootstrap simulations, where bootstrap samples are drawn from the inferred distribution of projected summary statistics as function of the parameters (see Supplementary Section S.1.7 for details about this procedure). Various definitions exist for bootstrap-based confidence intervals (e.g., Davison and Hinkley, 1997, Chapter 5). We thus compare the coverage of intervals defined by inverting profile-LRTs whose p-values are read from the *χ*^2^ distribution, to two bootstrap intervals. One interval is defined by inverting the profile-LRTs whose p-values are read from the bootstrap distribution of the likelihood-ratio statistic. The second bootstrap interval uses a Bartlett correction (Bartlett, 1937): it is defined by inverting p-values read from the *χ*^2^ distribution, for the profile likelihood ratio statistic corrected by the estimated mean of this statistic in the bootstrap samples. The definitions of the three confidence intervals are summarized in Table 1 and their coverage values are reported in later Tables as “profLR”, “bootLR” and “BcorCI” respectively. We also considered percentile bootstrap intervals, but these appear to have less predictable coverage over the different simulations conditions.

**Table 1:**
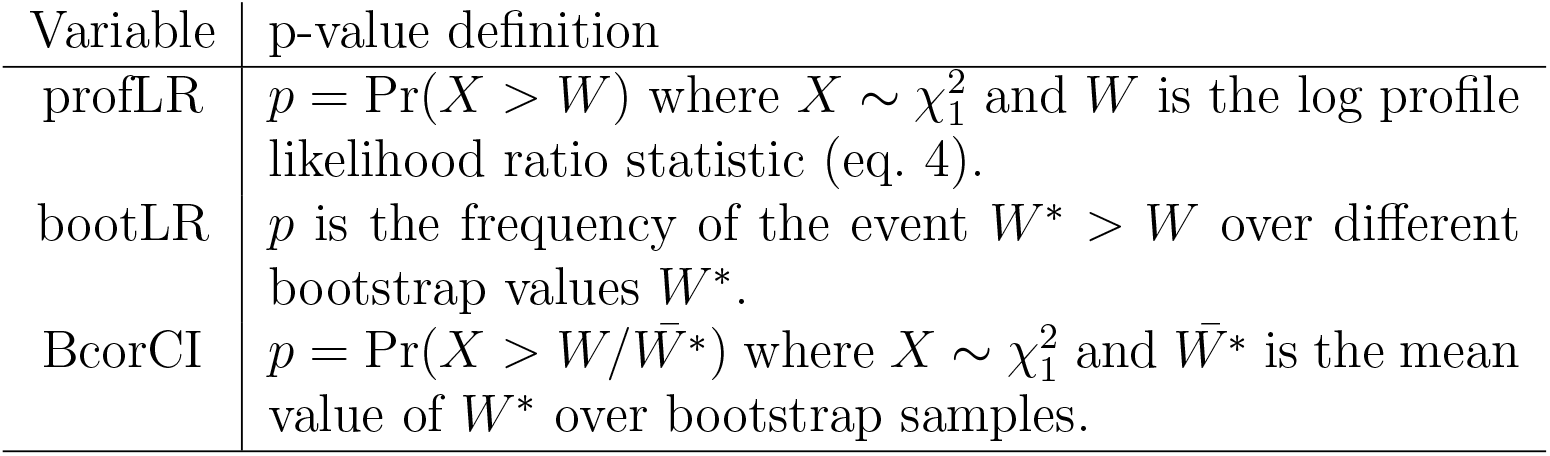
Tests that yield p-values and profile LRTs used to define confidence intervals. *W*^∗^ is the log profile likelihood ratio statistic (eq. 4) evaluated on a parametric bootstrap sample *S*^∗^ from the fitted model. Realized coverage of implied confidence intervals at nominal level *C* is the frequency of the event *p >* 1 − *C* over simulated data sets.

We report the conditional coverage of two types of credibility intervals: the central posterior intervals computed by the `abcrf` R package (Marin *et al*., 2022), and highest posterior density (HPD) intervals also deduced from the ABC-RF output.

It is not always reasonable to require exact control of the coverage or distribution of p-values. If there is no information about a parameter in the data (i.e., if likelihood is flat with respect to it), the conclusion of the analysis should be that there is no information. This lack of information is represented by unbounded intervals with 100% coverage. On the other hand, weak information will result in intervals with more than the nominal coverage and in distributions of summary-LRTs that show a deficit of low p-values relative to the uniform distribution. These patterns readily occur in our simulations of demo-genetic scenarios.

## Results

### 15-parameter toy model

In this model, we consider the estimation of the covariance matrix of a 5-dimensional multivariate normal distribution. It has two variants, one where all 15 parameters are identifiable, the other where only 5 variances are identifiable. As this example mainly serves as a first check of the inference workflow in a case with a relatively high number of parameters and where we know which parameters are identifiable and which are not, no variation of data-generating parameters nor any comparison with ABC are presented.

Performance summaries are detailed in Supplementary Table S.2 and distributions of p-values of summary-LRTs are shown in Fig. 2. In each case 400 datasets were simulated and reference tables of 44000 samples were constructed independently for each dataset. Coverage is approximately controlled for identifiable parameters, mean coverage over the twenty rows of Table S.2 being 0.954 for the profile-likelihood ratio intervals and 0.96 for the Bartlett-corrected bootstrap intervals. There is expectedly a strong deficiency of low p-values for unidentifiable ones. Ideally, in the latter case, the distribution of p-values should be a step function in *p* = 1, and the observed non-stepwise distributions are the result of the fluctuations of the estimated surface compared to the exact one, these fluctuations being too small to result in low p-values. Overall, when parameters are not identifiable, the confidence intervals still have at least nominal coverage.

**Figure 2:**
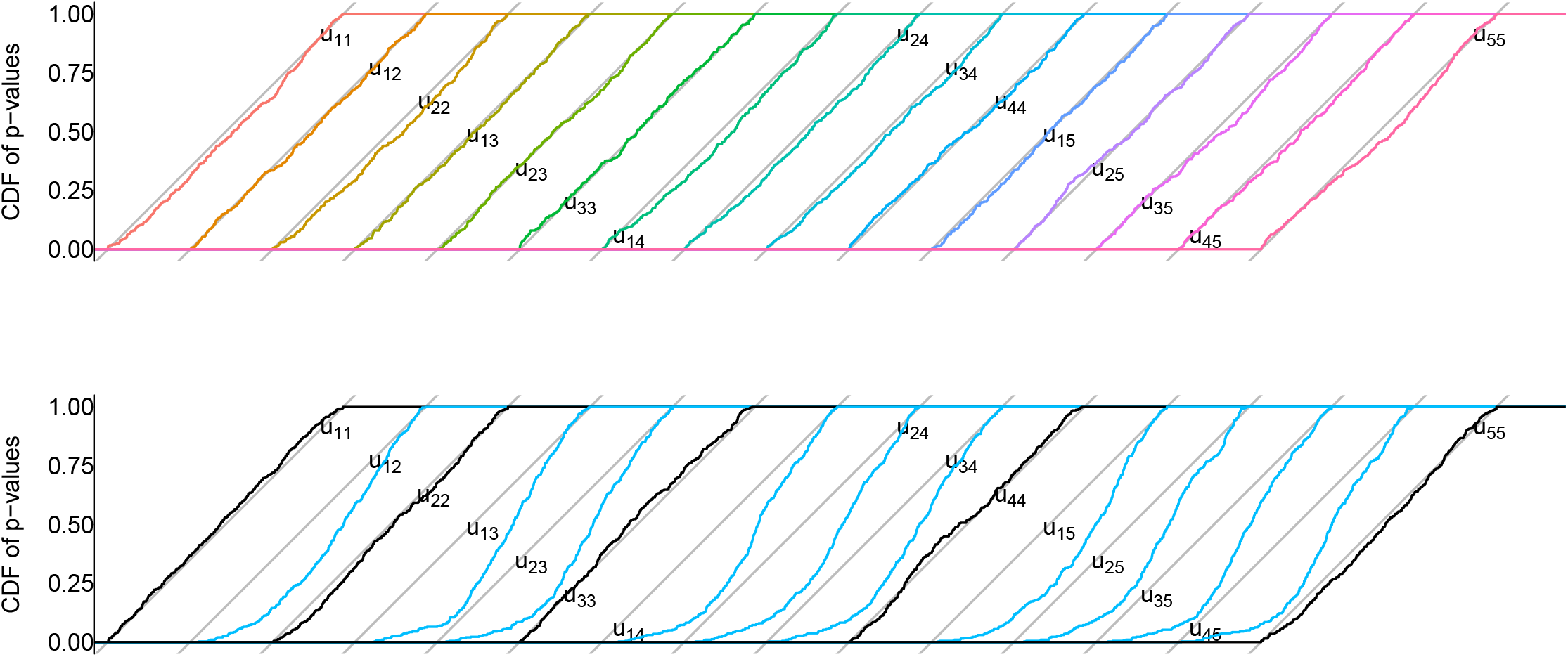
Distributions of p-values of summary-LRTs in multivariate-normal toy example. The cumulative distributions of p-values are shown for the tests of each parameter of the covariance matrix (i.e., each element *u*_*ij*_ of its Cholesky factor, eq. S.2 in Supplementary Material). The grey diagonal lines represent uniform densities on [0,1], shifted in the x axis for visibility. Top: fully-identifiable model. Bottom: partially-identifiable model, with distributions for identifiable and non-identifiable parameters shown in black and blue, respectively. The distributions for non-identifiable parameters show a marked deficiency of low p-values (the cumulative distribution being under the diagonal), as expected since the inferred likelihood surface should be flat with respect to such parameters.

### Ladybird invasion scenario (8 parameters)

The simulation results reported in Table 2 show that there is little information about admixture rate *r*_a_. Posterior estimators for this parameter appear to have low bias only because the data-generating value 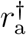 of *r*_a_ is close to the mean value of the prior distribution. This also explains why these posterior estimators have lower RMSE (conditional on 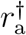) than the summary-MLE. There is also weak information about 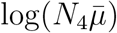, *t*_1_, and 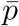. Coverage of the likelihood-ratio intervals is too high, 98% on average, for these four parameters, but also for the four other ones (97,05% on average, Table 2, “profLR” column). These excesses largely disappear in the bootstrap-corrected versions of the summary-LRTs, even for the parameters with low information (Table 2, “bootLR” and “BcorCI” columns, and Supplementary Fig. S.1, bottom), with the Bartlett-corrected intervals having 95.4% coverage on average for the four better-estimated parameters and 95.5% for the four other ones.

**Table 2:** Performance summaries for the ladybird invasion scenario. Bias and root-mean-square error (RMSE) are reported for the maximum-summary likelihood estimates (MSLE), and the posterior mean (postEv) and posterior median (postMed) for ABC-RF. Coverage of confidence intervals with nominal 95% level is reported for summary-likelihood inference (profLR) as implied by the distribution of p-values shown in Fig.S.1, for two forms of bootstrap correction described in the Text (profLR and BcorCI), for the central intervals provided by ABC-RF (postCI), and for highest posterior density intervals (HPD CI). Bold font is used to emphasize for each parameter the bias value minimal in absolute value, the minimum RMSE, and the coverage closest to 0.95.

| parameter | Relative bias |  |  | Relative RMSE |  |  | coverage |  |  |  |  |
| --- | --- | --- | --- | --- | --- | --- | --- | --- | --- | --- | --- |
|  | MSLE | postEv | postMed | MSLE | postEv | postMed | profLR | bootLR | BcorCI | postCI | HPD CI |
| $\log(N_1\bar{\mu})$ | **0.0008** | 0.010 | 0.008 | **0.035** | 0.036 | 0.036 | 0.970 | 0.965 | 0.967 | **0.960** | 0.997 |
| $\log(N_2\bar{\mu})$ | **-0.0004** | -0.078 | -0.064 | **0.092** | 0.148 | 0.150 | 0.975 | **0.950** | 0.945 | 0.980 | 0.975 |
| $\log(N_3\bar{\mu})$ | **0.009** | -0.040 | -0.024 | **0.096** | 0.129 | 0.134 | 0.965 | **0.952** | 0.960 | 0.990 | 0.990 |
| $\log(N_4\bar{\mu})$ | 0.044 | **0.026** | 0.056 | 0.264 | **0.050** | 0.073 | 0.992 | **0.952** | 0.960 | 1.000 | 1.000 |
| $t_1$ | **0.049** | 0.128 | 0.081 | 0.281 | 0.149 | **0.131** | 0.987 | 0.940 | **0.947** | 1.000 | 1.000 |
| $r_a$ | 0.017 | 0.013 | **0.008** | 0.171 | **0.103** | 0.121 | 0.972 | 0.960 | **0.957** | 1.000 | 1.000 |
| $\log(\bar{\mu})$ | **-0.012** | -0.100 | -0.078 | **0.064** | 0.129 | 0.111 | 0.972 | **0.952** | 0.945 | 0.985 | 0.982 |
| $\bar{p}$ | **0.006** | 0.023 | 0.018 | 0.171 | **0.143** | 0.146 | 0.967 | **0.952** | **0.947** | 0.987 | 0.985 |

The ABC-RF estimators of the different parameters often have higher bias, but there is no clear trend for RMSEs, consistently with the fact that the relative magnitude of the RMSE of ML vs. posterior estimates depends on where the data-generating values lie within the prior support. While the dependence of relative RMSE performance on location of the data-generating parameters in the prior distribution may explain some of the heterogeneity in apparent performance of the two methods, additional simulations show that more obscure specificities of the ABC-RF method contribute to this heterogeneity. Notably, the ABC-RF estimates for 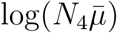 are biased away from the prior mean (−0.5), their correlation with summary-ML estimates is low (0.215), and they have a much lower variance than the latter. Together, these results are not expected from general theory for posterior estimators, given the uniform priors and the fact that summary-MLE estimates do not exhibit a similar bias. Varying the 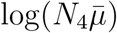 value, with other parameters being fixed, shows that the ABC-RF 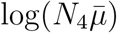 estimator is biased away from the prior mean over much of the prior range (Supplementary Fig. S.2), whereas a bias toward the prior mean would be more commonly expected (e.g., Casella and Berger, 2002, Section 7.3.4). Yet, it has lower averaged RMSE (Supplementary Table S.6) than the summary-ML estimator, as expected when comparing the prior-averaged performance of ML and posterior-mean estimators.

In Section S.4.2 of the Supplementary Information, we assessed performance of inferences for alternative data-generating parameter values derived from a preliminary fit of the actual data. These simulations exhibit even less information about parameters *t*_1_ and *r*_a_, and repeat the striking patterns of Fig. S.2 for estimation of 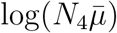.

Overall, this means that ABC-RF estimators can be strongly biased toward a value quite distinct from the prior mean, and their variance around this value may be small. ABC-RF estimates may then exhibit much higher or much lower RMSE than summary-MLEs, depending on the data-generating parameter values ***θ***^*†*^ chosen, and this pattern cannot be simply interpreted in terms of location of ***θ***^*†*^ relative to the prior mean.

### 7-parameter human admixture scenario

This inference scenario is obtained by fixing the following ten parameters from the full 17-parameter scenario schematized in the right part of Fig. 1 : log(*N*_1_) = 4.1, log(*N*_3_) = 4.3, log(*N*_4_) = 3.5, *r*_a_ = 0.2, *d*_3_ = 42, *N*_bn3_ = 160, *d*_4_ = 9, *N*_bn4_ = 98, log(*N*_34_) = 3.1, *d*_34_ = 24.

Performance summaries are reported in Table 3 and Supplementary Fig. S.4. The summary-likelihood method generally exhibits lower bias and RMSE than ABC-RF (Table 3). ABC-RF estimates can be strongly biased. In particular, we observe for parameter *t*_23_ a bias away from the prior mean, similarly to the pattern previously discussed for log(*N*_4_) in the ladybird invasion scenario. In a transformed log scale, the data-generating value was log(1 + *t*_23_) = 1.5, the mean summary-MLE was 1.66, the realized prior mean in the ABC-RF reference table was 1.855 and the mean ABCRF estimate was 2.491. Examination of the log-likelihood values *ℓ*(***θ***) for parameters values ***θ*** from both the ABC-RF reference table and from the summary-likelihood reference table for a given dataset suggests an explanation, which we illustrate in Fig. 3, left panel (similar results have been obtained for other simulated datasets). All *ℓ*(***θ***) are here the values of the log-likelihood function estimated by the summary-likelihood inference for the dataset. The points from the ABC-RF reference table form a cloud with a likelihood maximum near *x* = 2.5. This readily explains the mean posterior estimate near 2.5. In this case, the uniform prior sampling used by the ABC-RF method hence appears to miss the more relevant parameter region. One consequence is that the conditional coverage of posterior intervals for *t*_23_ is quite low (15.25%). On the other hand, the iterative exploration of the parameter space by the summary-likelihood workflow allows higher likelihood values to be attained for lower *t*_23_ values.

**Table 3:** Performance summaries for the 7-parameter human admixture scenario. Bias and root-mean-square error (RMSE) are reported for the maximum-summary likelihood estimates (MSLE), and the posterior mean (postEv) and posterior median (postMed) for ABC-RF. Coverage of confidence intervals with nominal 95% level is reported for summary-likelihood inference (profLR), for two forms of bootstrap correction described in the Text (bootLR and BcorCI), for the central intervals provided by ABC-RF (postCI), and for highest posterior density intervals (HPD CI). Bold font is used to emphasize for each parameter the bias value minimal in absolute value, the minimum RMSE, and the coverage closest to 0.95.

| parameter | Relative bias |  |  | Relative RMSE |  |  | coverage |  |  |  |  |
| --- | --- | --- | --- | --- | --- | --- | --- | --- | --- | --- | --- |
|  | MSLE | postEv | postMed | MSLE | postEv | postMed | proflR | bootLR | BcorCI | postCI | HPD CI |
| $\log(N_2)$ | **0.007** | 0.100 | 0.105 | **0.047** | 0.103 | 0.108 | 0.955 | **0.947** | **0.947** | **0.952** | 0.980 |
| $t_1$ | **0.101** | 0.304 | 0.293 | 0.317 | 0.313 | **0.309** | 0.985 | 0.967 | **0.965** | 1.000 | 0.997 |
| $\log(t_{12})$ | **-0.004** | -0.033 | -0.029 | **0.046** | 0.056 | 0.057 | 0.960 | **0.942** | 0.927 | 1.000 | 1.000 |
| $\log(1 + t_{23})$ | **0.039** | 0.245 | 0.275 | **0.069** | 0.246 | 0.276 | 0.972 | 0.970 | **0.950** | 0.152 | 0.195 |
| $N_{\text{bn}34}$ | **0.007** | 0.359 | 0.326 | **0.019** | 0.362 | 0.334 | 0.992 | 0.980 | 0.982 | 0.885 | **0.955** |
| $\log(1 + t_{34})$ | 0.006 | -0.037 | **-0.004** | 0.028 | 0.046 | **0.022** | **0.947** | 0.937 | 0.940 | 1.000 | 1.000 |
| $\log(N_a)$ | **0.004** | 0.080 | 0.103 | **0.052** | 0.085 | 0.108 | 0.962 | 0.945 | **0.947** | 1.000 | 1.000 |

**Figure 3:**
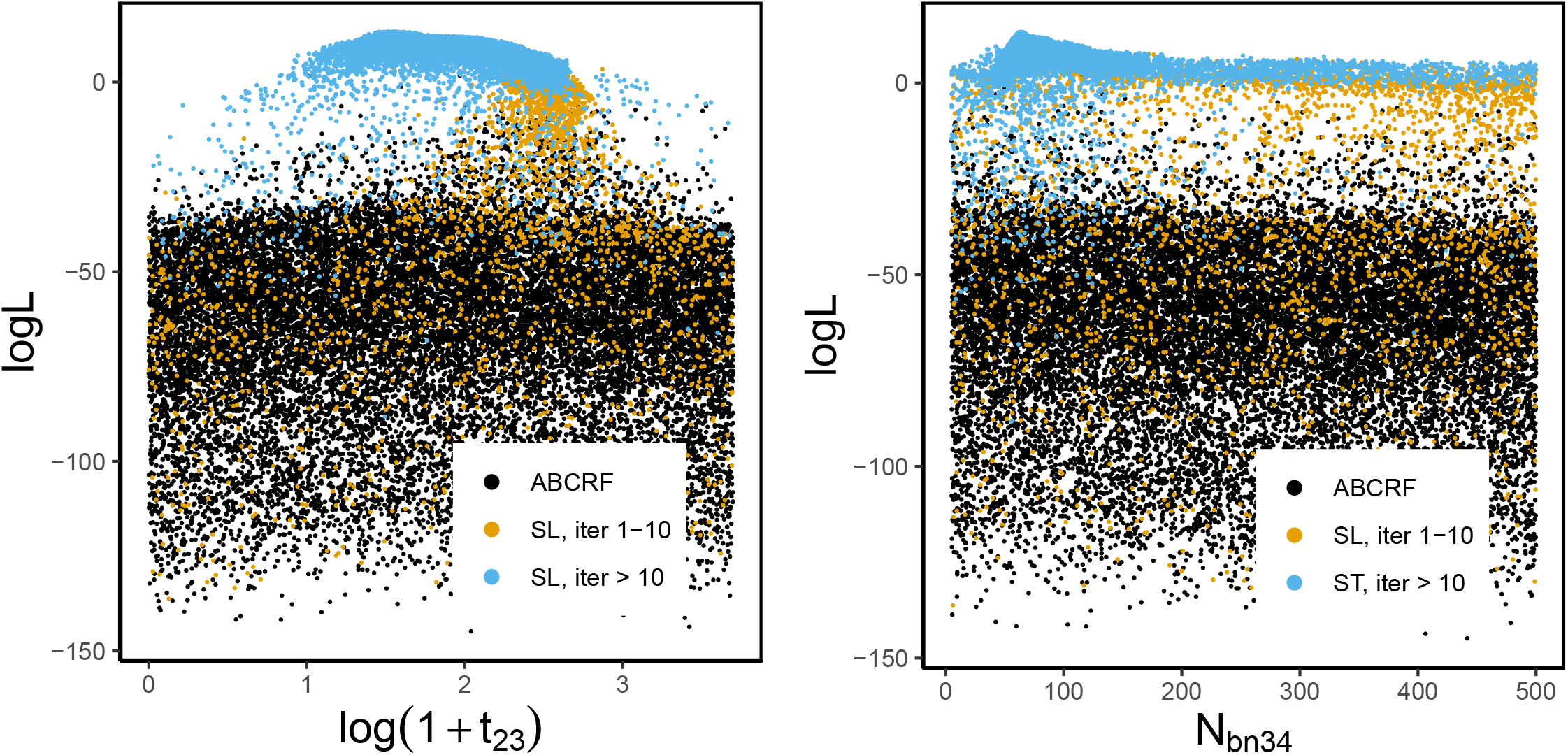
log-likelihoods of points ***θ*** from ABC-RF and summary likelihood reference tables. *x*-axis values are those of log(1+*t*_23_) (left) or *N*_bn34_ (right) from each ***θ***, for points from the ABC-RF reference table (in black) and for points from the summary inference reference table (in orange for points from the first 10 iterations, and in blue for points from later iterations). *y*-axis values are log-likelihood values according to the likelihood surface inferred in the final iteration of the summary likelihood workflow.

For both *t*_1_ and *N*_bn34_, estimates provided by ABC-RF are largely driven by the mean of the prior distribution, which is more expected in the case of a parameter with less apparent information about it (*t*_1_) than with more information about it (*N*_bn34_). The latter case might be explained as a less striking effect of the failure of the ABC-RF analysis to explore the narrow region of parameter space with high likelihood (Fig. 3, right panel). Two consequences of this failure, combined with attraction of posterior estimates towards the prior mean, are that the conditional coverage of “central” posterior intervals for *N*_bn34_ is distinctively low (88.5%), but that HPD intervals are less affected.

Deficiencies of low p-values are observed for LRTs, without bootstrap correction, for admixture time *t*_1_, bottleneck size *N*_bn34_, and time *t*_23_, with corresponding higher than nominal coverage of the intervals. For *t*_1_ there is little statistical information in the data: likelihood profiles are rather flat, and many estimates are at the lower bound of the parameter space. For *N*_bn34_ there is more information, as shown in particular by the low RMSE of summary-MLEs. The coverage of the Bartlett-corrected intervals for the other parameters is 94.6% on average.

To mitigate the effects of a low level of information in the data for some parameters, we performed additional simulations with datasets of 10,000 SNP rather than 5,000 as previously assumed. The detailed results reported in Supplementary Table S.11 show reductions of RMSE of summary-MLEs by a factor between 1.36 and 1.58 for the different parameters except *N*_bn34_, which exhibits a reduction by 2.02. By contrast, the RMSE of the posterior mean estimator is reduced by a lower factor (between 0.93 and 1.28). Hence, the summary-likelihood results exhibit reductions in RMSE roughly as expected from doubling the information in the data (i.e. a reduction of RMSE by a 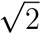 factor), but ABC-RF does not. This could reflect another benefit of a more efficient exploration of the region of interest of the parameter space by the iterative workflow. Indeed, when larger datasets are considered, the parameter region with high likelihood ratios becomes narrower and is thus expected to become more difficult to appraise by non-iterative methods. We thus expect iterative methods to provide larger gains in precision relative to non-iterative ones for larger datasets.

We have repeated the simulations as in Table 3 except that the *N*_bn34_ value was set to either the lower or the upper bound of its explored range (this parameter was chosen because it is the best estimated among the seven parameters, in terms of relative RMSE). The results, detailed in Supplementary Table S.12, show that, as expected, HPD intervals perform much better than central intervals in this case, where they perform similarly to profile-likelihood based intervals.

### 13-parameter Human admixture scenario

Results for the 13-parameter simulation scenario are presented in Table 4 and Supplementary Fig. S.6, with composite bottleneck intensity parameters defined as ratios of bottleneck duration to bottleneck population size: *b*_3_ = *d*_3_*/N*_bn3_, *b*_4_ = *d*_4_*/N*_bn4_, and *b*_34_ = *d*_34_*/N*_bn34_. Summary-MLEs have lower absolute bias than posterior estimates, while for RMSEs the pattern is more heterogeneous, and can be interpreted as resulting from a combination of the effects previously considered to explain discrepancies between the two methods. A more detailed analysis for each parameter is presented in Supplementary Section S.5.3.2. In particular, it suggests that poor exploration of parameter space again affects estimation of log(1 + *t*_23_), but also of log(*t*_12_), log(*N*_bn3_), log(*N*_bn4_), and *b*_34_. The admixture rate *r*_a_ and the composite bottleneck intensity parameters *b*_3_ and *b*_4_ appear well estimated by both methods, and population size *N*_2_ may also be relatively well estimated by SL.

**Table 4:** Performance summaries for the 13-parameter human admixture scenario. Bias and root-mean-square error (RMSE) are reported for the maximum-summary likelihood estimates (MSLE), and the posterior mean (postEv) and posterior median (postMed) for ABC-RF. Coverage of confidence intervals with nominal 95% level is reported for summary-likelihood inference (profLR), for two forms of bootstrap correction described in the Text (bootLR and BcorCI), for the central intervals provided by ABC-RF (postCI), and for highest posterior density intervals (HPD CI). Bold font is used to emphasize for each parameter the bias value minimal in absolute value, the minimum RMSE, and the coverage closest to 0.95.

| parameter | Relative bias |  |  | Relative RMSE |  |  | coverage |  |  |  |  |
| --- | --- | --- | --- | --- | --- | --- | --- | --- | --- | --- | --- |
|  | MSLE | postEv | postMed | MSLE | postEv | postMed | proflR | bootLR | BcorCI | postCI | HPD CI |
| $\log(N_1)$ | **-0.121** | -0.296 | -0.279 | 0.363 | 0.302 | **0.290** | 0.967 | **0.947** | 0.955 | 1.000 | 1.000 |
| $\log(N_2)$ | **-0.006** | 0.027 | 0.023 | 0.033 | 0.034 | **0.031** | **0.962** | 0.935 | **0.937** | 1.000 | 1.000 |
| $t_1$ | **0.076** | 0.252 | 0.195 | 0.248 | 0.261 | **0.220** | 0.980 | 0.932 | **0.940** | 1.000 | 1.000 |
| $r_a$ | **-0.001** | -0.002 | -0.002 | **0.015** | 0.017 | 0.019 | **0.955** | 0.925 | 0.930 | 0.997 | 1.000 |
| $\log(t_{12})$ | **-0.020** | -0.055 | -0.061 | **0.056** | 0.066 | 0.077 | 0.997 | **0.992** | 0.995 | 1.000 | 1.000 |
| $d_3/N_{\text{bn}3}$ | **0.008** | 0.030 | 0.017 | 0.033 | 0.039 | **0.027** | 0.990 | **0.967** | **0.967** | 1.000 | 1.000 |
| $\log(N_{\text{bn}3})$ | **-0.075** | -0.133 | -0.146 | 0.164 | **0.138** | 0.152 | 1.000 | **0.972** | 0.982 | 1.000 | 1.000 |
| $d_4/N_{\text{bn}4}$ | **0.001** | 0.008 | -0.002 | 0.025 | **0.024** | 0.026 | 0.985 | **0.962** | 0.972 | 1.000 | 1.000 |
| $\log(N_{\text{bn}4})$ | **-0.021** | -0.069 | -0.086 | 0.185 | **0.081** | 0.109 | 0.992 | **0.965** | 0.975 | 1.000 | 1.000 |
| $\log(1+t_{23})$ | **0.022** | 0.204 | 0.234 | **0.106** | 0.206 | 0.236 | 0.985 | **0.967** | 0.970 | 1.000 | 1.000 |
| $d_{34}/N_{\text{bn}34}$ | **-0.033** | -0.109 | -0.108 | **0.055** | 0.111 | 0.110 | 0.982 | **0.972** | 0.982 | 1.000 | 1.000 |
| $\log(N_{\text{bn}34})$ | **0.002** | 0.017 | -0.035 | 0.121 | **0.029** | 0.045 | 0.985 | **0.960** | 0.967 | 1.000 | 1.000 |
| $\log(N_a)$ | **-0.009** | -0.037 | -0.018 | 0.042 | 0.045 | **0.038** | 0.965 | **0.940** | 0.935 | 1.000 | 1.000 |

Other parameters are poorly estimated by both methods. For log(*N*_1_) and *N*_bn34_ in particular, the position of the prior mean relative to the data-generating *θ*^*†*^ value, rather than efficient use of information, explains the low RMSE of the ABC-RF estimates. In Supplementary Section S.17, we show that summary-likelihood estimates are not improved by running further iterations of the workflow.

Intervals generally have higher than nominal coverage, which is not unexpected given the limited information present in the data about most parameters, and given additional boundary effects such as those detailed for *b*_3_ in Supplementary Section S.5.3.5. The coverage approaches the nominal 95% for the best estimated parameters, and the bootstrap corrections partially correct the coverage for the other parameters. These results may be seen as evidence that the summary-likelihood method is able to provide confidence intervals for clearly identifiable parameters, from limited simulation effort in such a 13-parameter scenario.

Finally, we compared the results of summary-likelihood and ABC-RF inferences from the real SNP dataset used to design our simulation-based study (Supplementary Table S.16). It is generally not easy to make sense of such comparisons unless additional empirical information, not included in the data, is available about the values to be inferred. We found that the estimates ≈ 0.2 of the admixture rate *r*_a_ (the proportion of genes of European ancestry within African American individuals) reported by both methods are consistent with previous studies (as discussed by Collin *et al*., 2021). The most notable pattern was that for the four parameters identified by the simulation study as being estimable with some precision, *r*_a_, *b*_3_, *b*_4_ and *N*_2_, the confidence intervals of our summary-likelihood method lay within the credible intervals of the ABC-RF method, and were two to five times narrower than the latter intervals. The largest ratio is observed for *N*_2_, which was found in the simulation to be much better estimated by summary-likelihood than by ABC-RF.

### Comparison to SNLE

We aimed to compare the performance of the summary likelihood approach with SNLE across the four simulation scenarios. However, we did not include SNLE for the 13-parameter admixture example because the parameter space is subject to non-rectangular constraints (Eq. S.3), which the `sbi` implementation does not readily accommodate. Addressing these constraints within the `sbi` framework would require at least a non-trivial reparameterization of the parameters. In the remaining scenarios, point-estimate performance (bias and RMSE) was broadly similar between the two methods, in particular when contrasted to ABC-RF. In contrast, interval estimation yielded more heterogeneous results (Table 5). In the 15-parameter toy example, SNLE posterior intervals achieved good coverage, comparable to summary-likelihood, but were consistently slightly wider (width ratio 1.02–1.16 across parameters; mean 1.07). By contrast, in the 7-parameter admixture scenario, SNLE intervals were mostly too narrow and exhibited poorly calibrated coverage. Results for the ladybird invasion scenario were intermediate: coverage was, on average, improved relative to uncorrected profile likelihood-ratio intervals, but showed greater variability across parameters (0.912–1).

**Table 5:** Compared performance of the summary likelihood and SNLE approaches. Bias and root-mean-square error (RMSE) are reported for the maximum-summary likelihood estimates (MSLE), and the posterior mean (postEv) for SNLE. Coverage of intervals with nominal 95% level is reported for summary-likelihood inference (profLR) for the central posterior intervals provided by SNLE (postCI). Boldface indicates, for each parameter, the smallest absolute bias, the minimum RMSE, and the coverage closest to 0.95.

| parameter | Summary likelihood |  |  |  |  | SNLE |  |  |
| --- | --- | --- | --- | --- | --- | --- | --- | --- |
|  | bias | RMSE | profLR | bootLR | BcorCI | bias | RMSE | postCI |
| 15-parameter toy example |  |  |  |  |  |  |  |  |
| $u_{11}$ | **0.003** | **0.048** | 0.970 | 0.970 | **0.967** | 0.027 | 0.056 | 0.925 |
| $u_{12}$ | **-0.003** | **0.071** | **0.945** | 0.962 | 0.960 | -0.021 | 0.075 | 0.930 |
| $u_{22}$ | **-0.001** | **0.063** | 0.967 | 0.980 | 0.970 | 0.026 | 0.073 | **0.960** |
| $u_{13}$ | **0.006** | **0.063** | 0.962 | 0.975 | 0.965 | 0.010 | 0.069 | **0.947** |
| $u_{23}$ | **-0.001** | **0.062** | 0.962 | 0.982 | 0.970 | -0.003 | 0.066 | **0.952** |
| $u_{33}$ | **-0.008** | **0.069** | 0.922 | 0.942 | 0.940 | 0.020 | 0.076 | **0.945** |
| $u_{14}$ | **0.002** | **0.063** | 0.947 | 0.972 | 0.970 | 0.018 | 0.069 | **0.950** |
| $u_{24}$ | **0.0009** | **0.058** | **0.952** | 0.960 | 0.955 | -0.011 | 0.063 | 0.940 |
| $u_{34}$ | **-0.003** | **0.050** | 0.955 | 0.965 | **0.952** | 0.008 | 0.053 | 0.955 |
| $u_{44}$ | -0.014 | **0.050** | 0.935 | **0.952** | 0.945 | **-0.003** | 0.051 | 0.957 |
| $u_{15}$ | **0.006** | 0.060 | 0.970 | 0.977 | 0.972 | 0.020 | **0.057** | **0.950** |
| $u_{25}$ | **0.0009** | **0.052** | **0.952** | 0.967 | 0.960 | 0.002 | 0.055 | **0.947** |
| $u_{35}$ | -0.001 | **0.050** | **0.952** | 0.965 | 0.960 | **-0.0003** | 0.052 | 0.955 |
| $u_{45}$ | 0.006 | **0.046** | **0.947** | 0.960 | 0.942 | **0.001** | 0.048 | **0.947** |
| $u_{55}$ | -0.018 | 0.050 | 0.932 | **0.945** | **0.945** | **-0.009** | **0.049** | **0.955** |
| Ladybird invasion scenario |  |  |  |  |  |  |  |  |
| $\log(N_1\bar{\mu})$ | 0.0008 | **0.035** | 0.970 | **0.965** | 0.967 | **-0.00008** | 0.036 | 0.912 |
| $\log(N_2\bar{\mu})$ | **-0.0004** | 0.092 | 0.975 | **0.950** | 0.945 | -0.009 | **0.081** | 0.930 |
| $\log(N_3\bar{\mu})$ | 0.009 | 0.096 | 0.962 | **0.952** | 0.960 | **0.002** | **0.083** | 0.927 |
| $\log(N_4\bar{\mu})$ | **0.044** | 0.264 | 0.992 | **0.952** | 0.960 | 0.077 | **0.124** | 0.977 |
| $t_1$ | **0.049** | 0.281 | 0.987 | 0.940 | **0.947** | 0.105 | **0.174** | 1.000 |
| $r_a$ | **0.017** | 0.171 | 0.972 | 0.960 | **0.957** | 0.017 | **0.146** | 0.962 |
| $\log(\bar{\mu})$ | **-0.012** | 0.064 | 0.972 | **0.952** | 0.945 | -0.023 | **0.063** | 0.942 |
| $\bar{p}$ | **0.006** | 0.171 | 0.967 | **0.952** | **0.947** | 0.016 | **0.151** | **0.947** |
| 7-parameter Human admixture scenario |  |  |  |  |  |  |  |  |
| $\log(N_2)$ | 0.007 | 0.047 | 0.955 | **0.947** | **0.947** | **0.002** | **0.043** | 0.845 |
| $t_1$ | **0.101** | 0.317 | 0.982 | 0.967 | 0.965 | 0.211 | **0.270** | **0.937** |
| $\log(t_{12})$ | **-0.004** | 0.046 | 0.960 | **0.942** | 0.927 | -0.009 | **0.042** | 0.820 |
| $\log(1 + t_{23})$ | 0.039 | 0.069 | 0.972 | 0.970 | **0.950** | **0.032** | **0.057** | 0.775 |
| $N_{bn34}$ | **0.007** | **0.019** | 0.997 | 0.980 | 0.982 | 0.009 | 0.028 | **0.977** |
| $\log(1 + t_{34})$ | 0.006 | 0.028 | **0.947** | 0.937 | 0.940 | **0.005** | **0.026** | 0.890 |
| $\log(N_a)$ | 0.004 | 0.052 | 0.960 | 0.945 | **0.947** | **-0.0001** | **0.046** | 0.880 |

## Discussion

In this study, we present and assess the performance of an automated workflow for summary-likelihood, a method of simulation-based inference based on inferring a likelihood surface for summaries of the data. The efficiency of the method is contingent upon its iterative procedure of exploration of the parameter space. In a 15-parameter toy example, the coverage of confidence intervals provided by summary-likelihood is nearly optimal. However, in cases where some parameters are not practically identifiable, the results are inherently more complex. In these cases, the intervals derived from the likelihood profiles for parameters with low information content are, as expected, too large. Nevertheless, bootstrap procedures appear to be a viable solution for obtaining better coverage. Our simulations suggest that the “bootLR” bootstrap intervals may be systematically used to improve the coverage of intervals. These intervals are defined by the profile-likelihood ratio threshold value determined as the *q*-quantile of the bootstrap distribution of the likelihood ratio, where *q* is the intended coverage. The Bartlett-corrected intervals often performed similarly to these intervals. We also considered bootstrap-corrected intervals based on the distribution of parameter estimates, namely the “basic” and “percentile” intervals (Davison and Hinkley, 1997). However, their coverage (only reported in the electronic Supplementary Material) was more variable across simulation conditions, so we cannot recommend them as a general-purpose option.

Although we could not perform an extensive simulation study of the sensitiveness of the performance of our workflow to its control parameters, we performed simulation of the effect of increasing of decreasing the number of Gaussian components in MGMs (Supplementary Section S.3.2.2). We also verified that increasing the size of the reference table from 38000 to 50000 in the 13-parameter admixture model yields no clear benefit (Supplementary Section S.17). Conversely, we found that decreasing this size from 20000 to 6000 in the 7-parameter model led to degraded performance (Supplementary Table S.15).

Beyond documenting the general performance of the summary-likelihood method, our simulations provide a comparison with the ABC-RF method. Raynal *et al*. (2019) compared their ABC-RF method to some iterative (or “sequential”) ABC workflows (ABC-PMC, Beaumont *et al*., 2009; Prangle, 2017; SMC-ABC, Del Moral *et al*., 2012). For a toy example, they found ABC-RF to better infer marginal posterior distributions, without the computational overhead of the iterative methods. However, they did not extend their comparisons to their population genetics example. By contrast, our results highlight the benefits of an iterative workflow for the exploration of parameter space, and are thus consistent with the premises of sequential ABC methods.

There are further differences between the summary-likelihood and ABC-RF methods, beyond the nature of the intervals sought, and whether the workflow is iterative or not. In particular, ABC-RF is currently constrained to infer marginal posterior distributions for each parameter separately, which makes it more difficult to identify sets of parameters that can be estimated only in combination with each other. By contrast, the likelihood surface inherently retains information about the joint effect of parameters on the likelihood of the data. It is also possible to extend ABC-RF for the inference of multivariate parameters, using distributional random forests (Dinh *et al*., 2024).

The comparisons with ABC-RF are based on previously considered simulation scenarios. Our findings indicate that ABC-RF is capable of producing estimates with favorable bias and RMSE characteristics, comparable to those of summary-likelihood. However, we also identified instances where its estimation performance diverged significantly, which can be attributed, at least in part, to the imperfect exploration of parameter space by non-iterative methods. Further evidence of this imperfect exploration can be observed in the markedly smaller reduction in the root mean square error (RMSE) of ABC-RF estimates relative to SL estimates when the amount of data is increased, as was investigated in the 7-parameter human admixture scenario.

Moreover, we confirmed the previous observation that ABC-RF often produces 95% credibility intervals with 100% coverage. This is true even for the most easily estimated parameter. For instance, Raynal *et al*. (2019, Fig. 2) already found that 95% credibility intervals for the admixture rate *r*_a_ in the Human admixture scenario had 100% coverage. Comparison of credibility and confidence intervals shows that in such cases the credibility intervals consistently extend in both directions beyond the confidence intervals.

A further issue is that ABC-RF estimates may be significantly biased. This phenomenon can be attributed to the fact that sampling from pre-specified priors, a common practice in non-sequential ABC methods, is inadequate for fully exploring the parameter space in certain models. Consequently, the credibility interval coverage provided by ABC-RF may be considerably diminished in such instances.

We found that replacing reference tables, each constructed for a different simulated dataset, with a single reference table of the same size leads to poor inference, even if it is obtained by subsampling the reference tables constructed for each different simulated dataset (Supplementary Section S.5.2.4). While this result has little impact on the analysis of real datasets, each requiring a new reference table matching the details of the sampling design of the dataset, it prevents a drastic reduction of computation time in our simulation studies, which would be possible if a single reference table could be used for accurate inference from all simulated datasets.

Random-forest regression has been used in this work to reduce the dimension of the summary statistics. The implementation of our workflow allows other reduction methods to be used. However, we used the Random-forest regression approach here for reasons previously discussed (Collin *et al*., 2021; Pudlo *et al*., 2016; Raynal *et al*., 2019) and because it provides a convenient baseline for comparing other components of the inference workflow. This approach is fast, easily automated, and although it is not necessarily the most efficient, the additional effort needed to develop more efficient summaries on a case-by-case basis should be considered when designing applications of simulation-based inference. For example, Quelin *et al*. (2025) compared random forests, gradient boosting, and neural network methods. Neural networks achieved a 5.8% reduction in RMSE compared to random forests (average value other the 10 cases in their Tables 1 and 2), for 10 to 100 fold increases in training times (their supplementary Figure S1). Moreover, a preliminary study was necessary to optimize their alternative methods separately for each inferred parameter.

In our simulations, the size of the reference table was determined only by the number of parameters. Our implementation allows alternative and more adaptive stopping criterion to be used, based on the estimated precision of the profile likelihood ratios at the bounds of the inferred confidence intervals. A bootstrap procedure is implemented in our R package to evaluate mean square errors of prediction of these ratios. It is thus possible to request termination of iterations when the average RMSE for the different bounds, or when all RMSEs, are below a certain threshold. However, such a termination condition is necessary rather than sufficient, because likelihood surfaces may be inferred with high precision but low accuracy, in particular in cases where large reference tables are built from few iterations, whereas many iterations would be needed to identify narrow parameter regions with high likelihood.

The fixed reference-table sizes in our simulations were deliberately kept small enough to allow performance evaluation over hundreds of samples, but larger tables may be required to achieve accurate inference in many applications. In particular, the linear scaling with the number of parameters used here may become increasingly optimistic as dimensionality grows. Further, even with few parameters, much larger numbers of simulations may be required when the distribution of the data is heavy-tailed, as illustrated by the g-and-k distribution (Frazier *et al*., 2024).

Further iterative ABC methods have been developed, notably sequential neural likelihood estimation (SNLE, Papamakarios *et al*., 2019) and related workflows based on deep-learning methods that learn probability distributions and related functions (e.g., Sharrock *et al*., 2024 and references therein). In our comparison of summary-likelihood with SNLE, we found that the relative performance of the two methods strongly depended on the simulation scenario. SNLE showed excellent performance in one case but more variable interval coverage in the other two, particularly in the 7-parameter admixture scenario. As currently implemented, the main limitation of summary-likelihood is its rapidly increasing runtime with the number of parameters, whereas SNLE’s iterative likelihood learning becomes comparatively more efficient as dimensionality grows. This leaves open the possibility that a workflow based on iterative MAF training to learn the likelihood surface, as in SNLE, could ultimately deliver consistently better-calibrated intervals at more moderate computational costs.

## Conclusion

In this study, we have implemented and evaluated summary-likelihood inference, an iterative simulation-based method for approximate likelihood inference. In addition to documenting the method’s favorable performance, including in regard to confidence interval inference, we have conducted comparisons with the ABC-RF method. These comparisons demonstrate that constructing a reference table of simulations based on pre-specified priors may result in the omission of the most pertinent parameter regions. In contrast, an iterative method of exploring the parameter space may prove more effective in identifying these regions. Using summary-likelihood inference, we found that the full inference for any given dataset required a moderate number of simulations, of the order of 3000 times the number of estimated parameters, showing that approximate likelihood inference is a practically achievable objective even when intensive simulation of genomic datasets is required.

While iterative ABC methods have the potential to outperform non-iterative simulation-based approaches such as standard rejection ABC or ABC-RF, they are less commonly used by non-experts. This may be due to factors such as simplicity, robustness, computational requirements and familiarity with non-iterative approaches. However, as the field advances and tools become more user-friendly, there should be a shift towards iterative methods, particularly for studies involving high-dimensional parameter spaces and where computational efficiency is critical. The SNLE iterative method, which uses MAF training to learn the likelihood surface, already enables fast and efficient inference, but the intervals it provides do not always appear to be well-calibrated. Our method can yield better intervals within a reasonable time-frame for models with a moderate number of parameters, but our results leave open the possibility that improved use of iterative MAF training may also provide better intervals at even lower computational costs.

## Acknowledgments

We thank A. Courtiol, D. Prangle, P. Druilhet and an anonymous reviewer for comments on the manuscript, and the Genotoul bioinformatics platform Toulouse Occitanie (https://doi.org/10.15454/1.5572369328961167E12) for providing computing resources. This work has been supported by funds from the Occitanie Regional Council’s program “Key challenge BiodivOc” managed by the University of Montpellier (DevOCGen project), and has benefited from state aid managed by the project AgroStat (reference: ANR-23-EXMA-0002) of the Maths-VivES France 2030 program handled by the French Agence Nationale de la Recherche. A preprint version of this article has been peer-reviewed and recommended by PCI Math Comp Biol (https://doi.org/10.24072/pci.mcb.100426).

## Code and data availability

The source code of the `diyabc` simulator used in this study is available at https://github.com/diyabc/diyabc/releases/tag/v1.1.36. The R packages used in this study are all available on https://cran.r-project.org/. Additional data and scripts are available both at https://doi.org/10.5281/zenodo.19615138 and https://github.com/f-rousset/SI_InfusionMS.

## Supplementary Material

### S.1 Control parameters of algorithms

#### S.1.1 Iterative sampling of parameter points

The aim of the iterative procedure is to fill a target “top” space defined as the space of parameters with a given minimum likelihood ratio *l*_min_ relative to the summary-MLE. As a first approach, a uniform sampling of the currently predicted top space, together with sampling out of it with decreasing density for decreasing likelihood, may seem appropriate to identify iteratively the boundaries of the target space and to infer the likelihood surface within it. An approximately uniform sampling of the target space is obtained by a simple rejection sampling method, where parameter vectors are first sampled from a given proposal distribution *q*_**Θ**_ covering the target space, and the following rejection step preferentially retains sampled points with low *q*_**Θ**_ density, with probability in inverse proportion of the *q*_**Θ**_ density relative to its maximum among the sampled points. The current estimate *î*_**Θ**_ of the instrumental density is used as proposal distribution, except as specified below. To explore the boundaries, distinct rejection probabilities are also computed for points outside the top region. These probabilities not only depend on the *q*_**Θ**_ density but also on the likelihood ratios.

We have met some problems when applying such rules. From a broad initial instrumental distribution, few of the randomly sampled points have likelihood high enough to be retained after the rejection step, so fewer points may be retained than a given target number *n*_*i*+1_ unless an impractically large number of candidate points is sampled (in practice the number of candidate points generated has been bounded to essentially 10*n*_*i*+1_ even if the acceptance rate in previous iterations was much lower than 0.1). This problem is of particular concern in high-dimensional spaces. This can be improved by the use of the current instrumental posterior density *î*_**Θ**|*T*_ (. |**p**[**u**(𝒟)]) as a proposal distribution for the sampling of a fraction of the points. More precisely, the current *î*_**Θ**_ is used to obtain 90% of the points (as a target: maybe less if rejection rate is high), and the instrumental posterior is used to generate the missing points.

Another problem that was apparent, at least in the early versions of the workflow, is that these rules may sample too few points on the boundaries of the top space. Several modifications have been implemented to favor exploration at these boundaries. In the current workflow, performance may be practically insensitive to these modifications, except in the early iterations and for cases such as those where uniform sampling of parameter space leads to poor ABC-RF performance. Firstly, points are sampled not from each of the multivariate Gaussian distributions of the MGM model, but from the corresponding multivariate Student distributions (with dispersion matrix as given by the Gaussian covariance matrix, and degree of freedom set to 1). Secondly, the proportion of retained points at the boundary can be controlled by rescaling their rejection probabilities. A rescaling of the likelihood ratio of outside points sampled from *î*_**Θ**_ has been implemented so that the proportion of outside points among retained ones can increase to a given value *q*. All simulations used *q* = 1*/*4. The rejection rule for points sampled from the instrumental posterior distribution is more complex, but also designed to sample points at the boundaries of the top space.

These rules depart from the idea of sampling from the posterior distribution, applied in deep-learning based iterative workflows. However, for uniform priors, sampling new parameters from the posterior distribution is equivalent to sampling with relative probabilities in proportion to likelihood ratios, and is then similar to a simple variant of our sampling procedure. We implemented such sampling in proportion to likelihood ratios, still with rejection sampling, and still allowing fewer points to be retained than the pre-specified target number. This means that iterations still proceed by smaller steps when rejection sampling has low success rate. We compared the two versions in two test cases (Tables S.14 and S.10). With the current workflow and its default controls, the performance of the two sampling algorithms are indistinguishable. Only by focusing on parameters for which uniform sampling of parameter space leads to distinctly poor ABC-RF performance, and on the early iterations, can we detect some effect of the sampling rule (Table S.15). For the default sampling rule, this Table also illustrates the effect of reducing the size of the reference table to 6000 samples, by comparison with Table 3 which reports results using 20000 samples, the recommended minimum for such a 7-parameter case.

#### S.1.2 Selection of simulated samples for projections

In each iteration *i*, a first subset *S*_*i*_ of points can be defined as follows. For each parameter point in the reference table, the predicted (summary-) log-likelihood is computed. Then all points with log-likelihood above 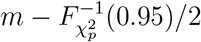 are retained, where *m* is the maximum predicted log-likelihood for the table, *p* is the number of estimated parameters, and 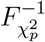 is the quantile function of the chi-squared distribution with *p* degrees of freedom. This criterion defines a parameter region analogous to a p-dimensional 95% confidence region, except that *m* is not the inferred maximum of the likelihood surface. The final subset for iteration *i* is the union of *S*_*i*_, of *S*_*i*−1_ (as using only *S*_*i*_ sometimes resulted in a form of periodic behavior of the projections and likelihood inferences over successive iterations), and of the initial reference table so as to allow training over the entire parameter ranges initially considered. When these rules select less than 1000*p* points, additional points from the top are retained up to that number.

#### S.1.3 Control of random-forest methods

Default values are defined for control settings of random-forest methods in `Infusion` and `abcrf`, which were used in all simulations reported below. 1000 trees were used in all cases for both methods. The other default settings differ between the two methods, as those from `Infusion` combine features of Breiman’s original random-forest method and of the extremely-randomized trees method (Geurts *et al*., 2006). The latter method does not provide out-of-bag predictions, but the combined options used as defaults by `Infusion` do provide such predictions. This combination is made possible by the `ranger` R package (Wright and Ziegler, 2017), and the combined defaults (`splitrule=`“`extratrees`”, `mtry` set to the number of predictor variables, `replace` set to `FALSE`, and `sample.fraction` set to 0.632) were chosen in light of the results of an independent simulation study of the effects of the settings on prediction performance (Appleton *et al*., 2022, Appendix), giving some slight advantage to these combined settings over the basic random-forest method.

#### S.1.4 Summary-likelihood maximization

Practical issues in the maximization of the inferred likelihood surface are the possible occurrence of local maxima, and premature termination of maximization procedures at a boundary of the parameter space. Such occurrences may become manifest when computing likelihood profiles and summary-LRTs, where it may appear that the inferred likelihood maximum was in fact only a local maximum, exceeded by the profile likelihood for some other parameter values. Although the issue of local maxima does not have a perfect solution, it is possible to draw many candidate vectors ***θ*** from the “instrumental posterior” distribution *î*_**Θ**|*T*_ (.|**p**[**u**(𝒟)]), derived from the current joint density estimate 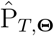 by conditioning on *T* = **p**[**u**(𝒟)]. This posterior distribution is easily deduced from the joint MGM model for the parameters and the projected summaries. Using the candidate vector ***θ*** with the highest predicted likelihood as the initial value for maximization appears to be relatively efficient in reducing the impact of local maxima on the final inference.

An additional problem of maximization steps is that a spurious maximum of the likelihood function may be inferred by extrapolation in poorly sampled parameter regions. To avoid this, a penalization has to be defined for likelihood predictions in such regions. This penalization compares the predicted parameter density of the parameter point whose likelihood is to be evaluated, to a threshold defined from the distribution of fitted parameter densities for sampled points at the top of the likelihood surface.

#### S.1.5 Distribution modelling

We fit a MGM model for the probability distribution of points **y** (here in joint parameters and projected statistics space), of the form

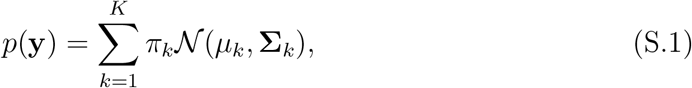

where *K* is the number of components, *π*_*k*_ is the probability that a point **y** belongs to component *k*, and 𝒩 (***µ***_*k*_, **Σ**_*k*_) is the multivariate Gaussian distribution with mean ***µ***_*k*_ and covariance matrix **Σ**_*k*_. For a given number of components, all parameters are jointly estimated by maximizing the likelihood without constraints on parameter space (especially without any enforced relationship between the different covariance matrices).

The number of components *K* must be selected. Akaike’s information criterion (AIC) may be used to select the number of components, which seems appropriate given that the retained model is used for prediction of modelled density. Other model selection criteria are often considered in the MGM literature, but for different purposes, such as estimating a “true” number of components of the mixture. For *n* points (here, the number of sample simulations in the reference table), Lebret *et al*. (2015) suggest trying *K* up to *n*^0.3^ components, a rule originating from Wong (1982) (see Bozdogan, 1994, p. 77, for discussion). For small-dimensional distributions, preliminary simulations showed that higher values, accessible by a minor increase of the exponent, may improve the summary-likelihood inference for large *n*. However, when applying such rules for given *n* and increasing dimension of the inferred distribution, the number *P* of parameters of the MGM model will exceed *n*, highlighting that a lower bound must be considered. So we defined two constraints on *K*: it is bounded by the constraint *P* ≤ *n/*4, and by ⌈*n*^0.31^⌉ . Given such rules for the maximum number of components, we found that for large reference tables (in terms of number of simulations times number of variables), the largest value is generally retained according to the AIC criterion. Thus, it is practically equivalent, but much less time-consuming than trying all possible numbers of components, to systematically set *K* to the upper bound defined by the above two constraints. This simple rule was applied systematically beyond the first iteration.

By application of these rules, for a given number *n* of samples in the reference table, *K* is effectively the largest number ≤ ⌈*n*^0.31^⌉ such that the number *P* of parameters of the MGM model (a function of *K* and of the dimension of **y**) satisfies *P* ≤ *n/*4. Variants of these two rules for *K* are assessed in Section S.3.2.2.

Given these controls, the other controls of the MGM fitting algorithms were set to their default values in the `Rmixmod` package, except that the maximum number of iterations of the Expectation-Maximization algorithm was increased to 1000, and that non-default initialization values may be used. The MGM fit may occasionally fail or return inappropriate results, such as a mixture containing a Gaussian component with vanishing variances. In such cases, the model is refitted using one of the alternative initialization methods implemented in `Rmixmod`, and if this still fails, the number of components of the mixture is reduced.

#### S.1.6 Control of iterative refinements

A target number *n*_*I*_ of samples controls the number of samples *n*_*i*_ added in any iteration *i*, i.e., between each MGM fit of the reference table. *n*_*i*_ may sometimes be lower than *n*_*I*_ depending on realized success frequency in rejection sampling. *n*_I_ varied between 500 and 625 for the different simulations in this work. The main drawback of increasing the number of iterations by reducing *n*_I_ for a given final number of points is the increasing computation time due to the MGM modelling performed in each iteration (with a significant increase for large dimensional distributions only) and to projection steps (in many if not all iterations, see Section Refinements of likelihood surface inference through iterations in the Main Text). Typical computation times are presented in Table S.1.

The number of points added in the first iterations are controlled in a distinct way from later ones because it is better to sample comparatively fewer points per iteration when the likelihood surface is yet relatively poorly estimated. We used at most *n*_1_ = 1000 in the first iteration, then added samples over several iterations by steps of *n*_*i*_ samples increasing essentially in proportion to powers of two over these iterations.

For example, for 7 estimated parameters, the first reference table contains 400 samples, then successive target numbers of added samples in the next 5 iterations are 62, 62, 125, 250, 501 to reach a total of 1400 samples. The target *n*_*i*_ in further iterations is 500, and iterations are performed until the reference table contains 20,000 samples. In practice, users do not have to set such values, which are default values set by the software as function of the number of estimated parameters.

Ideally, iterations should be repeated until a certain degree of accuracy is reached for the inference of the likelihood surface. A bootstrap procedure based on resampling of the MGM distributions is implemented to estimate the uncertainty in the inference of the likelihood surface, as quantified by the mean square error of the likelihood ratio at the inferred confidence bounds. This allows automatic termination of the iterations when a certain precision is reached. However, in the present simulation-based study the aim is rather to quantify the precision of inferences for a given size of the reference table, comparable to the size of reference tables used in an ABC inference. The final size was therefore fixed regardless of the achieved precision.

**Table S.1:** Approximate fit times (in seconds). These are the elapsed (“wall-clock”) times reported by the R code profiler, for fitting a single dataset using the default workflow design, with details for the three main components: sample simulation by `diyabc` simulator, random-forest analyses by `ranger` R package, and MGM modelling by `Rmixmod` R package. The first two steps used the parallelization procedures implemented in C++ in each software, with 10 CPU cores. The analyzed datasets in these fits were either the real data for the scenarios of historical demography (the variant with composite bottleneck parameters for the 13-parameter case), or the first simulated dataset of the toy multivariate normal example. All computations were run on a virtual Linux machine, on a Dell Precision 3581 laptop. GPU was not used for SNLE computations, as this was not faster, consistently with the `sbi` documentation.

| scenario | parameters | Summary-likelihood |  |  |  | SNLE |
| --- | --- | --- | --- | --- | --- | --- |
|  |  | total | diyabc | ranger | Rmixmod |  |
| Human admixture | 7 | 2118 | 610 | 900 | 576 | 1533 |
| Ladybird invasion | 8 | 2815 | 60 | 1217 | 1504 | 985 |
| Human admixture | 13 | 13691 | 1241 | 7555 | 4728 | — |
| toy Normal | 15 | 5949 | NA | NA | 5838 | 896 |

#### S.1.7 Simplified bootstrap procedure

We used the parametric bootstrap to simulate the distribution of the summary-likelihood ratio statistic and to deduce from it a corrected p-value. We used it also to compute bootstrap confidence intervals. In its canonical version, the parametric bootstrap would be too time-consuming, as it would imply repeating the full inference workflow on many simulated datasets. However, in the present case, we used two approximations. First, it is possible to obtain the parametric bootstrap samples directly as a vector of projected summary statistics, denoted **t**^∗^, since the distribution of such samples, given the parameter estimates as in standard parametric bootstrap, is a MGM distribution easily deduced from the representation of the joint distribution stored in the fit object. Second, we did not reconstruct references tables for each of these bootstrap samples. Instead, inferences are performed on **t**^∗^, using the available estimated joint and marginal distributions to evaluate the terms in the expression for the summary-likelihood of the bootstrap sample, *L*(***θ***; **t**^∗^) = P_*T*,**Θ**_(**t**^∗^, ***θ***)*/*P_**Θ**_(***θ***).

Only 199 bootstrap samples were used for each bootstrap computations. It is advised to use more samples when analyzing a given dataset. However, such a low number is enough to appreciate the correction brought by the bootstrap procedure, and there is no point in obtaining very accurate tests and intervals from many bootstrap samples for each simulated dataset, when coverage is evaluated from 200 or 400 datasets.

### S.2 Choice of data-generating parameter values, *θ*^†^, in demographic scenarios

We have used an automatic rule to select ***θ***^*†*^ values in a relatively simple way that is both unaffected by our knowledge of the position of ***θ***^*†*^ within the prior distributions used in ABC analyses, and more consistent with the real data than uniform sampling in the prior would be.

In practice, we first derived ***θ***^*†*^ values from preliminary summary-likelihood fits of actual data, with one adjustment. In the Human admixture scenario, we set the admixture time *t*_1_ to 6 generations, ignoring its often unrealistically high (*>* 15 generations) estimate from actual data (the simulations indeed show that this parameter cannot be estimated with precision), and other ***θ***^*†*^ values were set to the constrained summary-MLEs given *t*_1_ = 6.

We have departed from this automatic rule when seeking to find conditions where parameters could be estimated with more precision, and to answer questions raised by a first set of simulation results. This approach is sufficient to facilitate an exploration and comparison of the performance of SL and ABC-RF across a range of conditions. In particular, none of the ***θ***^*†*^ values generated in this above-described way were in the lower 5% or upper 5% of the prior ranges. Therefore, in one case, a parameter value was set at either bound of its explored range.

### S.3 15-parameter toy example

#### S.3.1 Details of simulation scenarios

In this example, the parameter space is the space of 5 × 5 covariance matrices. For easy exploration of this parameter space, a transformation was applied to the parametric matrices (but not to the simulated statistics), so that the transformed space is unconstrained (Pinheiro and Bates, 1996): transformed parameter values are the elements of the upper-triangular Cholesky factor **U** of the covariance matrix **V** (**V** = **U**^⊤^**U**).

We defined a variant of this model where only 5 of the 15 parameters are identifiable, by performing simulations in which the covariance matrix is the cross-product of the matrix retaining only the diagonal elements of the 15-parameter Cholesky factor. In such a way, only diagonal covariance matrices are produced, and only the 5 standard deviation parameters (on diagonal positions of the Cholesky factor) are identifiable.

For each of these two variants, 400 simulated data were generated for the following transformed parameters, presented as the elements of the upper triangle of the Cholesky factor:

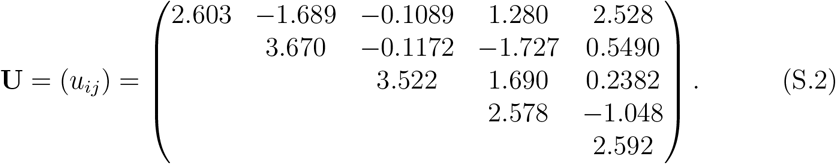

The corresponding data-generating covariance matrix **V**^*†*^ was obtained as a draw from a Wishart distribution with unit covariance matrix and 10 degrees of freedom. Parameters of the reference table were sampled in the ranges [0.5, 6] for the diagonal elements of the Cholesky factor, and [−4, 4] for the non-diagonal elements. These ranges were chosen to contain all values of Cholesky factors of 1000 samples from the same Wishart distribution as used to generate the above matrix.

#### S.3.2 Additional simulation results

##### S.3.2.1 Summary-likelihood inferences with default controls

**Table S.2:** Performance of inferences for multivariate Gaussian toy model. Last five lines are for the model where only the variances are identifiable. Bias and root-mean-square error (RMSE) are reported for the maximum-summary likelihood estimates. Coverage of confidence intervals with nominal 95% level is reported for summary-likelihood inference (profLR), and for two forms of bootstrap correction described in the Text (bootLR and BcorCI).

| parameter | rel. bias | rel. RMSE | profLR | bootLR | BcorCI |
| --- | --- | --- | --- | --- | --- |
| $u_{11}$ | 0.003 | 0.048 | 0.970 | 0.970 | **0.967** |
| $u_{12}$ | -0.003 | 0.071 | **0.947** | 0.962 | 0.960 |
| $u_{22}$ | -0.001 | 0.063 | **0.967** | 0.980 | 0.970 |
| $u_{13}$ | 0.006 | 0.063 | **0.962** | 0.975 | 0.965 |
| $u_{23}$ | -0.001 | 0.062 | **0.962** | 0.982 | 0.970 |
| $u_{33}$ | -0.008 | 0.069 | 0.920 | **0.942** | 0.940 |
| $u_{14}$ | 0.002 | 0.063 | **0.947** | 0.972 | 0.970 |
| $u_{24}$ | 0.0009 | 0.058 | **0.952** | 0.960 | 0.955 |
| $u_{34}$ | -0.003 | 0.050 | 0.955 | 0.965 | **0.952** |
| $u_{44}$ | -0.014 | 0.050 | 0.937 | **0.952** | 0.945 |
| $u_{15}$ | 0.006 | 0.060 | **0.972** | 0.977 | **0.972** |
| $u_{25}$ | 0.0009 | 0.052 | **0.952** | 0.967 | 0.960 |
| $u_{35}$ | -0.001 | 0.050 | **0.952** | 0.965 | 0.960 |
| $u_{45}$ | 0.006 | 0.046 | **0.945** | 0.960 | 0.942 |
| $u_{55}$ | -0.018 | 0.050 | 0.937 | **0.945** | **0.945** |
| $u_{11}$ | -0.001 | 0.051 | **0.947** | **0.947** | **0.947** |
| $u_{22}$ | 0.004 | 0.064 | **0.972** | 0.982 | 0.982 |
| $u_{33}$ | 0.003 | 0.069 | **0.952** | 0.955 | **0.952** |
| $u_{44}$ | -0.0006 | 0.049 | **0.957** | 0.962 | 0.960 |
| $u_{55}$ | 0.0007 | 0.045 | **0.965** | 0.972 | 0.972 |

##### S.3.2.2 Effect of alternatives rules for the number of clusters

As described in Section S.1.5, the number of clusters *K* of the multivariate Gaussian mixture model is the largest value ≤ ⌈*n*^0.31^⌉ and also such that *P* ≤ *n/*4, where *n* is the number of points, and *P* is the number of parameters of the mixture model.

The rule *P* ≤ *n/*4 reduces the maximum number of clusters in our 13- or 15-parameters inference scenarios. Table S.3 details the impact of the alternative rules *P* ≤ *n/*3 and *P* ≤ *n/*8 for the 15-parameter toy example. In this example, the final number of clusters are respectively 29, 22 and 11 when *P* ≤ *n/*3, *P* ≤ *n/*4 and *P* ≤ *n/*8 (for final size *n* = 44000 of reference table). The results for each of the 15 parameters considered individually do not clearly favor either rule. However, the coverage is almost always lower when *P* ≤ *n/*8, the average coverage being 0.930, versus 0.954 both when *P* ≤ *n/*3 and *P* ≤ *n/*4 (the latter rule being the default one, used in Table S.2 as in all other simulations).

**Table S.3:** Effect of alternatives to the *P* ≤ *n/*4 rule . These results are based on the same 400 simulated samples as in the first 15 rows of Table S.2. Bias and root-mean-square error (RMSE) of the maximum-summary likelihood estimates, and coverage of profile confidence intervals (profLR) with nominal 95% level, are reported for summary-likelihood inference. Minimal values of bias and RMSE, and coverage closest to 95% among the two compared rules, are emphasized in bold.

| parameter | $P \leq n/3$ | | | $P \leq n/8$ | | |
| --- | --- | --- | --- | --- | --- | --- |
|  | rel. bias | rel. RMSE | profLR | rel. bias | rel. RMSE | profLR |
| $u_{11}$ | 0.004 | **0.047** | 0.970 | **0.001** | 0.048 | **0.937** |
| $u_{12}$ | -0.004 | 0.071 | **0.947** | **-0.003** | **0.070** | 0.932 |
| $u_{22}$ | **-0.0001** | **0.064** | 0.972 | -0.003 | 0.065 | **0.945** |
| $u_{13}$ | 0.006 | **0.063** | 0.965 | **0.006** | 0.063 | **0.942** |
| $u_{23}$ | **-0.001** | 0.062 | 0.957 | -0.001 | **0.062** | **0.955** |
| $u_{33}$ | **-0.006** | **0.069** | **0.927** | -0.008 | 0.072 | 0.925 |
| $u_{14}$ | 0.003 | 0.063 | **0.952** | **0.002** | **0.063** | 0.925 |
| $u_{24}$ | **0.0007** | 0.059 | **0.955** | 0.001 | **0.058** | 0.920 |
| $u_{34}$ | **-0.003** | **0.049** | **0.952** | -0.005 | 0.050 | **0.947** |
| $u_{44}$ | -0.011 | **0.052** | **0.942** | **-0.008** | 0.057 | 0.937 |
| $u_{15}$ | 0.006 | 0.060 | **0.967** | **0.004** | **0.060** | **0.932** |
| $u_{25}$ | 0.0004 | 0.052 | **0.942** | **0.0003** | **0.052** | 0.922 |
| $u_{35}$ | -0.001 | 0.050 | 0.960 | **-0.001** | **0.049** | **0.942** |
| $u_{45}$ | **0.006** | 0.046 | 0.947 | 0.008 | **0.046** | **0.950** |
| $u_{55}$ | -0.011 | **0.055** | **0.952** | **-0.003** | 0.074 | 0.835 |

Conversely, when using the rule *P* ≤ *n/*4, the exponent of the power rule *K* ≤ ⌈*n*^0.31^⌉ only affects the number of clusters for relatively small *n*, i.e. in early iterations of the workflow, or for the final tables of the 7-parameter inference scenario. This scenario is not best suited for assessing the effect of the exponent, as coverage is affected by identifiability issues. Instead, we have simulated a 6-parameter toy multivariate Gaussian model defined exactly as the 15-parameter one, except that a 3 × 3 covariance matrix is considered instead of a 5 × 5 one. The summary statistics are thus defined as the 6 distinct elements of the observed covariance matrix of 50 draws from a multivariate normal distribution of dimension 3. The simulated data were drawn under the Gaussian distribution with the covariance matrix whose Cholesky factor is the 3 × 3 upper left block of the one shown in eq. S.2. Table S.4 compares the results of inferences using the exponents 0.29, 0.31 and 0.33, such that the number of clusters are respectively 17, 21 and 25 for the reference table of final size *n* = 17000 samples. The average coverages over the 6 parameters are respectively 0.9513, 0.9498 and 0.9480. The effect of the exponent is thus small, and the value 0.31 used in all other simulations appears adequate.

**Table S.4:** Effect of alternatives to the *n*^0.31^ rule. These results are based on 1000 simulated datasets. Bias and root-mean-square error (RMSE) of the maximum-summary likelihood estimates, and coverage of profile confidence intervals (profLR) with nominal 95% level, are reported for summary-likelihood inference. Minimal values of bias and RMSE, and coverage closest to 95% among the two compared rules, are emphasized in bold.

| parameter | $n^{0.29}$ | | | $n^{0.31}$ | | | $n^{0.33}$ | | |
| --- | --- | --- | --- | --- | --- | --- | --- | --- | --- |
|  | rel. bias | rel. RMSE | profLR | rel. bias | rel. RMSE | profLR | rel. bias | rel. RMSE | profLR |
| $u_{11}$ | 0.004 | 0.046 | **0.971** | 0.004 | 0.046 | 0.974 | **0.003** | **0.046** | 0.974 |
| $u_{12}$ | -0.006 | 0.068 | 0.960 | -0.006 | **0.068** | 0.956 | **-0.005** | 0.068 | **0.952** |
| $u_{22}$ | -0.001 | **0.067** | **0.950** | **-0.0009** | 0.067 | 0.948 | -0.001 | 0.067 | 0.945 |
| $u_{13}$ | -0.002 | **0.065** | 0.938 | **-0.002** | 0.065 | 0.936 | -0.002 | 0.065 | **0.940** |
| $u_{23}$ | 0.002 | **0.063** | **0.944** | **0.002** | 0.063 | 0.942 | 0.002 | 0.063 | 0.938 |
| $u_{33}$ | -0.006 | 0.063 | **0.945** | -0.005 | **0.063** | 0.943 | **-0.004** | 0.064 | 0.939 |

### S.4 Ladybird invasion scenario

#### S.4.1 Details of simulation scenarios

This scenario, described in Fig. 1 (left panel) of the Main Text, is derived from the study by Lombaert *et al*. (2011) on the worldwide invasion routes of the harlequin ladybird `Harmonia axyridis`, based on genotypes obtained at 18 microsatellite loci. More specifically, the scenario describes the genetic origin of the invasive West-European population (“Pop 4”; sample size: 32 diploid individuals) observed for the first time in 2001 and resulting from a genetic mixture between individuals originating from the invasive population of the Eastern USA (“Pop3”: 34 sampled individuals) and individuals originating from a European bio-control population (“Pop2”: 29 sampled individuals), the last two populations having been introduced from the species’ native area (i.e. China; “Pop1”: 29 sampled individuals) at known times (60 and 48 generations in the past for Pop2 and Pop3, respectively). A similar scenario was already taken as an example for ABC-RF analyses by Pudlo *et al*. (2016). The version studied here differs slightly from that of Lombaert *et al*. (2011), as the bottleneck events following each introduction are not modelled and are included in the effective sizes associated with each introduced population. On the other hand, the date of admixture *t*_1_ of the West-European population (Pop 4) is considered as a variable parameter of the model.

Sample simulations for this scenario have been run with version 1.1.36 of the `diyabc` command-line simulation program (Collin *et al*., 2021), with non-default `-o` argument allowing to provide the program with a table of sample-generating parameters. This allowed full control of the parameter values by the summary-likelihood procedures and avoided some boundary effects (see details in Supplementary Section S.5.1.1), which may otherwise occur when using `diyabc`.

In the reference tables used for our inferences, the raw microsatellite genotype data for the scenario studied are summarized by a total of 76 statistical summaries described in section 2.6.1 of the `diyabc` documentation (Collin *et al*., 2021; https://diyabc.github.io/doc/).

The assumed mutation model for the microsatellite markers is a generalized stepwise mutation model with two parameters, as implemented in `diyabc`: a geometric mean scale parameter 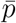 and a mean mutation probability parameter 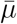. Both of these parameters control the distribution of locus-specific parameters: for each locus in each sample simulation, the locus-specific mutation parameter is drawn in a truncated gamma distribution with mean 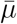 and shape 2, restricted to the range (10^−6^, 10^−2^); and the distribution of the number of repeats *X* of the microsatellite motif removed or added by a mutation event follows a geometric distribution *P* (*X* = *k*) = (1 − *p*)*p*^*k*−1^ where each locus-specific *p* is drawn from a truncated gamma distribution with mean 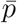 and shape 2, restricted to the range (0, 0.9).

The ranges of sampled parameter space were set to [−2, 1] for all 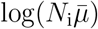, [3, 45] for *t*_1_, [0.05, 0.95] for *r*_a_, [−5, −2] for 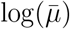, and [0.01, 0.05] for 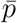.

**Table S.5:** The three different parameter vectors ***θ***^*†*^ used to generate simulated datasets in the 8-parameter examples of the ladybird invasion scenario. ^∗^: 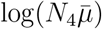 was variable among simulated datasets in Main Text Fig. S.2, and in supplementary Tables S.6, S.9 and Fig. S.3.

| $\log(N_1\bar{\mu})$ | $\log(N_2\bar{\mu})$ | $\log(N_3\bar{\mu})$ | $\log(N_4\bar{\mu})^*$ | $t_1$ | $r_a$ | $\log(\bar{\mu})$ | $\bar{p}$ | used in |
| --- | --- | --- | --- | --- | --- | --- | --- | --- |
| 0 | 0 | 0 | 0 | 13 | 0.5 | -3 | 0.25 | Table 2, Fig. S.1 |
| -0.031 | -0.84 | 0.56 | -0.88 | 19 | 0.5 | -2.7 | 0.22 | Tables S.7, S.10 |
| -0.031 | -1.5 | 0.75 | 0 | 9 | 0.2 | -2.7 | 0.22 | Tables S.8, S.9 |

##### S.4.1.1 Choice of data-generating parameter values, *θ*^*†*^

In principle, assessment of performance of confidence intervals calls for estimation of coverage for many different ***θ***^*†*^ values. However, this would be unpractical here due to the high-dimensional parameter space and the computational cost of the simulations. Instead, we have used an automatic rule to select ***θ***^*†*^ values in a relatively simple way that is both unaffected by our knowledge of the position of ***θ***^*†*^ within the prior distributions used in ABC analyses, and more consistent with the real data than uniform sampling in the prior would be.

In practice, we first derived ***θ***^*†*^ values from preliminary summary-likelihood fits of actual data, with one adjustment. In the Human admixture scenario, we set the admixture time *t*_1_ to 6 generations, ignoring its often unrealistically high (*>* 15 generations) estimate from actual data (the simulations indeed show that this parameter cannot be estimated with precision), and other ***θ***^*†*^ values were set to the constrained summary-MLEs given *t*_1_ = 6.

We have departed from this automatic rule when seeking to find conditions where parameters could be estimated with more precision, and to answer questions raised by a first set of simulation results. This approach is sufficient to facilitate an exploration and comparison of the performance of SL and ABC-RF across a range of conditions. However, none of the ***θ***^*†*^ values generated in this above-described way were in the lower 5% or upper 5% of the prior ranges. Therefore, in one case, a parameter value was set at either bound of its explored range.

The values of the parameters used to generate simulated data are shown in Table S.5 for the different studied cases.

#### S.4.2 Additional simulation results for the ladybird invasion scenario

##### S.4.2.1 Distribution of p-values for first set of parameter values

**Figure S.1:**
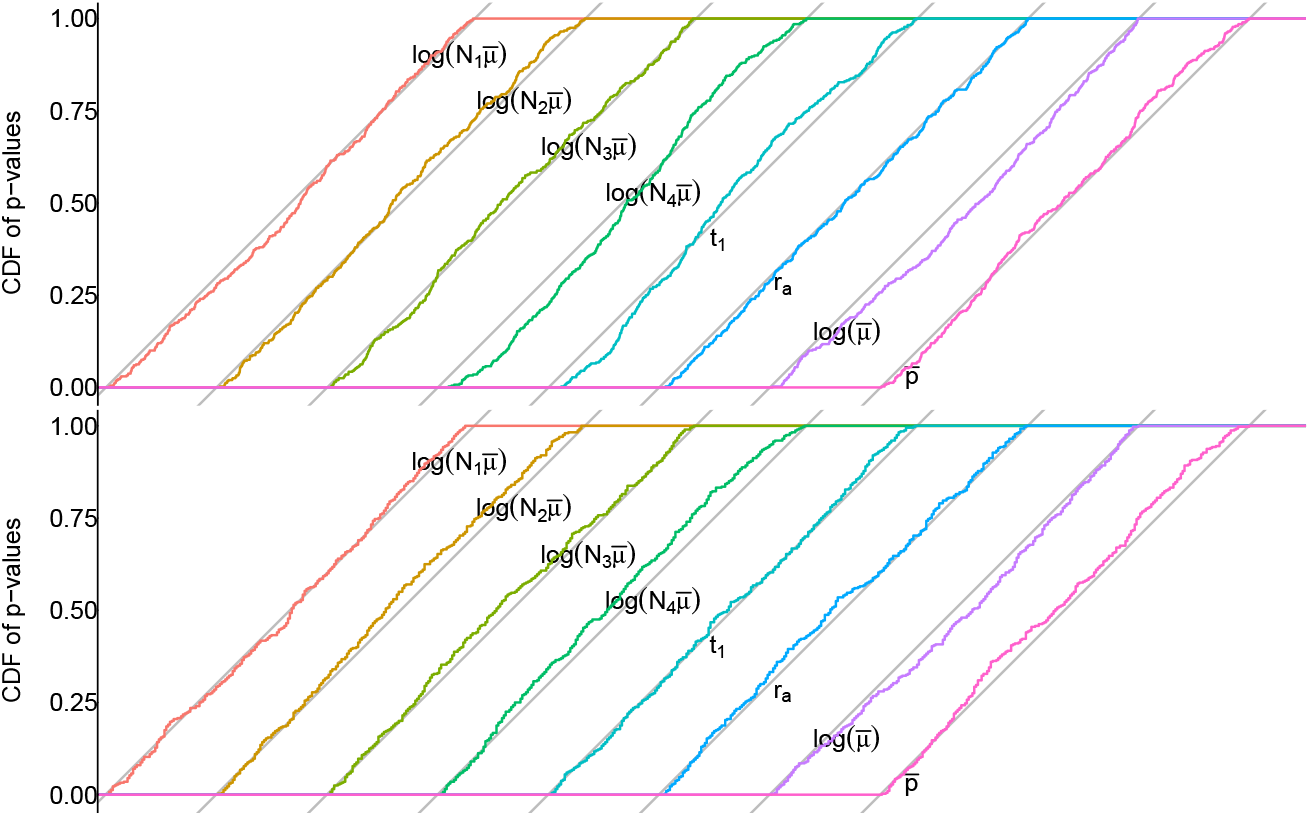
Cumulative distributions of p-values for the ladybird invasion scenario. This Figure describes the same simulation results as Main Text Table 2. The distributions of each uncorrected (top) and bootstrap-corrected (bottom) summary-LRTs (“bootLR” in Tables) for each parameter is shown relative to its own 1:1 reference line.

##### S.4.2.2 Performance summaries for variable 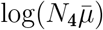

**Figure S.2:**
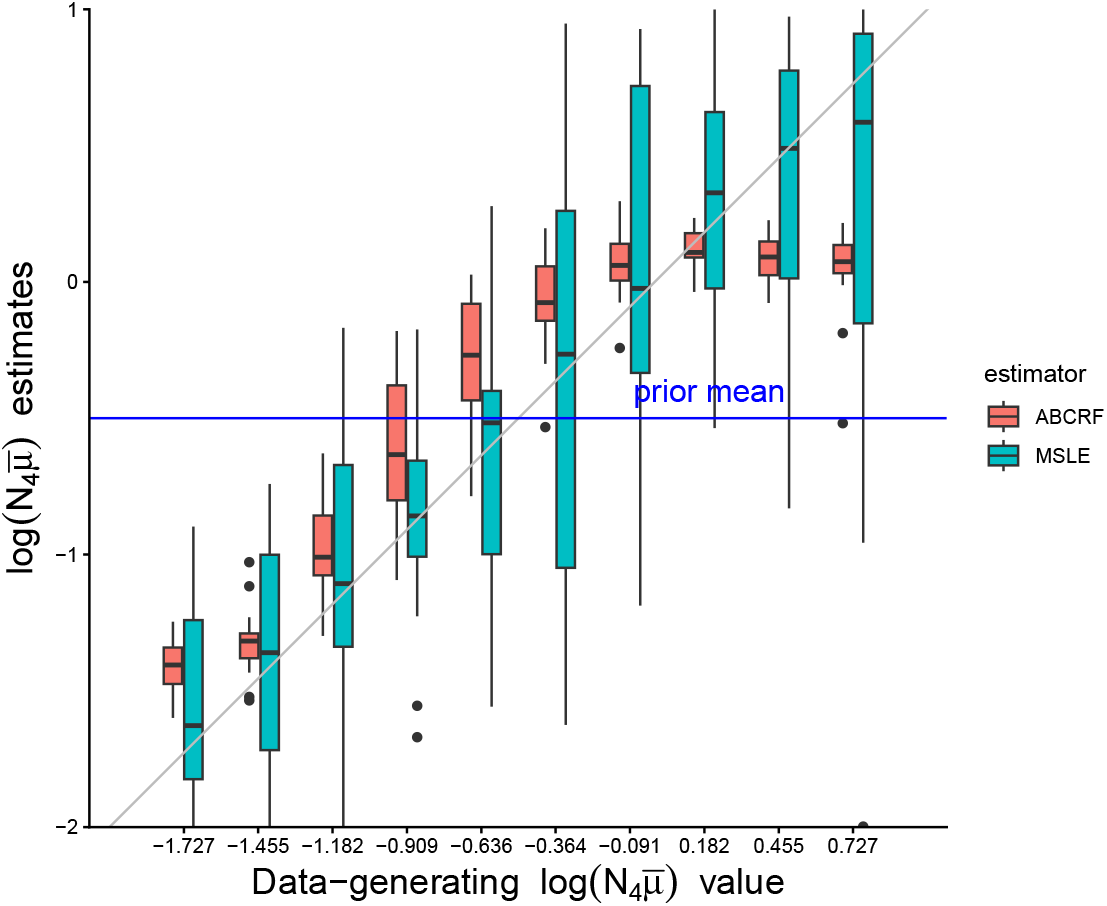
Distributions of MSLE and ABC-RF estimates of the parameter 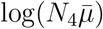. Twenty simulated samples were analyzed for each given 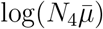 value. The first set of parameter values is used for the other parameters, as in Table 2.

**Table S.6:** Performance summaries for first set of parameter values and variable data-generating 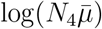. 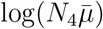 parameter values were varied as shown in Main Text Fig. S.2. 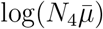 performance summaries are here average values over these variable 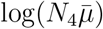 values. Other parameter values were constant as given by the first set of parameter values in Table S.5. Bias and root-mean-square error (RMSE) are reported for the maximum-summary likelihood estimates (MSLE), and the posterior mean (postEv) and posterior median (postMed) for ABC-RF. Coverage of confidence intervals with nominal 95% level is reported for summary-likelihood inference (profLR), for two forms of bootstrap correction described in the Text (bootLR and BcorCI), for the central intervals provided by ABC-RF (postCI), and for highest posterior density intervals (HPD CI). Bold font is used to emphasize for each parameter the bias value minimal in absolute value, the minimum RMSE, and the coverage closest to 0.95.).

| parameter | Relative bias |  |  | Relative RMSE |  |  | coverage |  |  |  |  |
| --- | --- | --- | --- | --- | --- | --- | --- | --- | --- | --- | --- |
|  | MSLE | postEv | postMed | MSLE | postEv | postMed | profLR | bootLR | BcorCI | postCI | HPD CI |
| $\log(N_1\bar{\mu})$ | **0.0004** | 0.009 | 0.008 | **0.035** | 0.036 | 0.036 | 0.975 | **0.945** | **0.945** | 0.960 | 0.995 |
| $\log(N_2\bar{\mu})$ | **0.004** | -0.039 | -0.028 | **0.090** | 0.120 | 0.123 | 0.980 | 0.965 | **0.955** | 0.990 | 0.985 |
| $\log(N_3\bar{\mu})$ | **0.004** | -0.028 | -0.015 | **0.093** | 0.118 | 0.125 | 0.980 | 0.965 | **0.950** | 0.995 | 0.995 |
| $\log(N_4\bar{\mu})$ | **0.007** | 0.021 | 0.037 | 0.191 | **0.121** | 0.122 | 0.985 | **0.945** | 0.965 | 1.000 | 1.000 |
| $t_1$ | **0.092** | 0.208 | 0.181 | 0.318 | 0.240 | **0.236** | 0.995 | **0.960** | 0.965 | 0.995 | 0.980 |
| $r_a$ | 0.012 | 0.014 | **0.010** | 0.248 | **0.107** | 0.135 | 0.975 | 0.955 | **0.950** | 1.000 | 1.000 |
| $\log(\bar{\mu})$ | **-0.014** | -0.078 | -0.059 | **0.063** | 0.111 | 0.096 | 0.980 | 0.960 | **0.955** | 0.990 | 0.985 |
| $\bar{p}$ | **0.019** | 0.030 | 0.023 | 0.199 | **0.157** | 0.160 | 0.935 | 0.935 | 0.925 | **0.960** | 0.930 |

In the following Tables we first present simulation results for two sets of data-generating parameter values differing from those of Table 2 in the Main Text, in order to check whether patterns in need of specific explanations persist in these different simulation conditions.

##### S.4.2.3 Second set of parameter values

Here, the data-generating values (Table S.5, second line) were chosen from a preliminary fit of the actual data. Nevertheless, the results (Table S.7) are overall similar to those of the Main Text, with even less information about parameters *t*_1_ and *r*_a_. 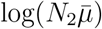 ABC-RF estimates are biased unexpectedly away from the prior mean. 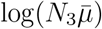, and to some extent *r*_a_ ABC-RF estimates, behave as those of low-information parameters, with a bias toward the prior. 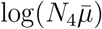 ABC-RF estimates have low bias, remarkably low variance and are uncorrelated to the SL estimates, as in the case from the Main Text.

**Table S.7:** Performance summaries for invasive ladybird scenario for second set of data-generating values. Bias and root-mean-square error (RMSE) are reported for the maximum-summary likelihood estimates (MSLE), and the posterior mean (postEv) and posterior median (postMed) for ABC-RF. Coverage of confidence intervals with nominal 95% level is reported for summary-likelihood inference (profLR), for two forms of bootstrap correction described in the Text (bootLR and BcorCI), for the central intervals provided by ABC-RF (postCI), and for highest posterior density intervals (HPD CI). Bold font is used to emphasize for each parameter the bias value minimal in absolute value, the minimum RMSE, and the coverage closest to 0.95.

| parameter | Relative bias |  |  | Relative RMSE |  |  | coverage |  |  |  |  |
| --- | --- | --- | --- | --- | --- | --- | --- | --- | --- | --- | --- |
|  | MSLE | postEv | postMed | MSLE | postEv | postMed | profLR | bootLR | BcorCI | postCI | HPD CI |
| $\log(N_1\bar{\mu})$ | **0.006** | 0.012 | 0.011 | **0.036** | 0.037 | 0.037 | 0.965 | 0.960 | **0.950** | 0.940 | 0.990 |
| $\log(N_2\bar{\mu})$ | **-0.007** | -0.069 | -0.063 | **0.049** | 0.085 | 0.086 | 0.965 | **0.945** | 0.930 | 1.000 | 1.000 |
| $\log(N_3\bar{\mu})$ | **0.019** | -0.050 | -0.030 | 0.099 | 0.087 | **0.086** | 0.980 | 0.940 | **0.950** | 1.000 | 1.000 |
| $\log(N_4\bar{\mu})$ | **-0.008** | -0.036 | -0.045 | 0.132 | **0.071** | 0.078 | 0.985 | **0.950** | 0.960 | 1.000 | 1.000 |
| $t_1$ | **0.065** | 0.150 | 0.157 | 0.285 | **0.160** | 0.172 | 1.000 | 0.960 | **0.950** | 1.000 | 1.000 |
| $r_a$ | **0.083** | 0.102 | 0.099 | 0.243 | **0.146** | 0.156 | 0.995 | **0.975** | 0.980 | 1.000 | 0.985 |
| $\log(\bar{\mu})$ | **-0.007** | -0.066 | -0.055 | **0.052** | 0.087 | 0.082 | 0.975 | **0.950** | 0.945 | 1.000 | 1.000 |
| $\bar{p}$ | **0.009** | 0.026 | 0.013 | 0.151 | **0.142** | 0.143 | 0.965 | 0.940 | **0.945** | 0.980 | 0.965 |

##### S.4.2.4 Third set of parameter values

This case is a variant of the previous one where the data-generating values for 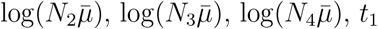 and *r*_a_ were modified (Table S.5, third line). The Results (Table S.8) are similar to those with the original data-generating values. In particular, the observations about 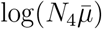 estimation are repeated. So we performed simulations for variable 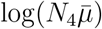 as in the Main Text, using the third set of values of other data-generating parameters. General patterns reported in the next Fig. S.3 and Table S.9 and are similar to those for the first set of simulation parameters (Table S.6, Main Text Fig. S.2).

**Table S.8:** Performance summaries for the invasive ladybird scenario for third set of parameter values. Bias and root-mean-square error (RMSE) are reported for the maximum-summary likelihood estimates (MSLE), and the posterior mean (postEv) and posterior median (postMed) for ABC-RF. Coverage of confidence intervals with nominal 95% level is reported for summary-likelihood inference (profLR), for two forms of bootstrap correction described in the Text (bootLR and BcorCI), for the central intervals provided by ABC-RF (postCI), and for highest posterior density intervals (HPD CI). Bold font is used to emphasize for each parameter the bias value minimal in absolute value, the minimum RMSE, and the coverage closest to 0.95.

| parameter | Relative bias |  |  | Relative RMSE |  |  | coverage |  |  |  |  |
| --- | --- | --- | --- | --- | --- | --- | --- | --- | --- | --- | --- |
|  | MSLE | postEv | postMed | MSLE | postEv | postMed | profLR | bootLR | BcorCI | postCI | HPD CI |
| $\log(N_1\bar{\mu})$ | **-0.002** | 0.006 | 0.005 | **0.035** | 0.035 | 0.035 | 0.975 | 0.955 | **0.950** | 0.980 | 0.995 |
| $\log(N_2\bar{\mu})$ | **-0.015** | -0.028 | -0.037 | 0.050 | **0.042** | 0.049 | 0.990 | 0.975 | **0.965** | 1.000 | 1.000 |
| $\log(N_3\bar{\mu})$ | **-0.009** | -0.123 | -0.106 | **0.089** | 0.135 | 0.120 | 0.985 | 0.965 | **0.955** | 1.000 | 1.000 |
| $\log(N_4\bar{\mu})$ | **0.040** | 0.057 | 0.075 | 0.263 | **0.061** | 0.078 | 0.995 | **0.970** | 0.990 | 1.000 | 1.000 |
| $t_1$ | **0.035** | 0.247 | 0.201 | **0.187** | 0.255 | 0.224 | 0.995 | **0.975** | 0.980 | 1.000 | 1.000 |
| $r_a$ | **0.011** | 0.057 | 0.027 | **0.044** | 0.075 | 0.053 | 0.965 | **0.940** | **0.940** | 1.000 | 1.000 |
| $\log(\bar{\mu})$ | **-0.014** | -0.043 | -0.038 | **0.051** | 0.062 | 0.060 | 0.960 | **0.945** | 0.940 | 1.000 | 1.000 |
| $\bar{p}$ | **0.013** | 0.028 | 0.019 | 0.153 | **0.140** | 0.143 | 0.975 | 0.955 | **0.950** | 0.985 | 0.975 |

**Figure S.3:**
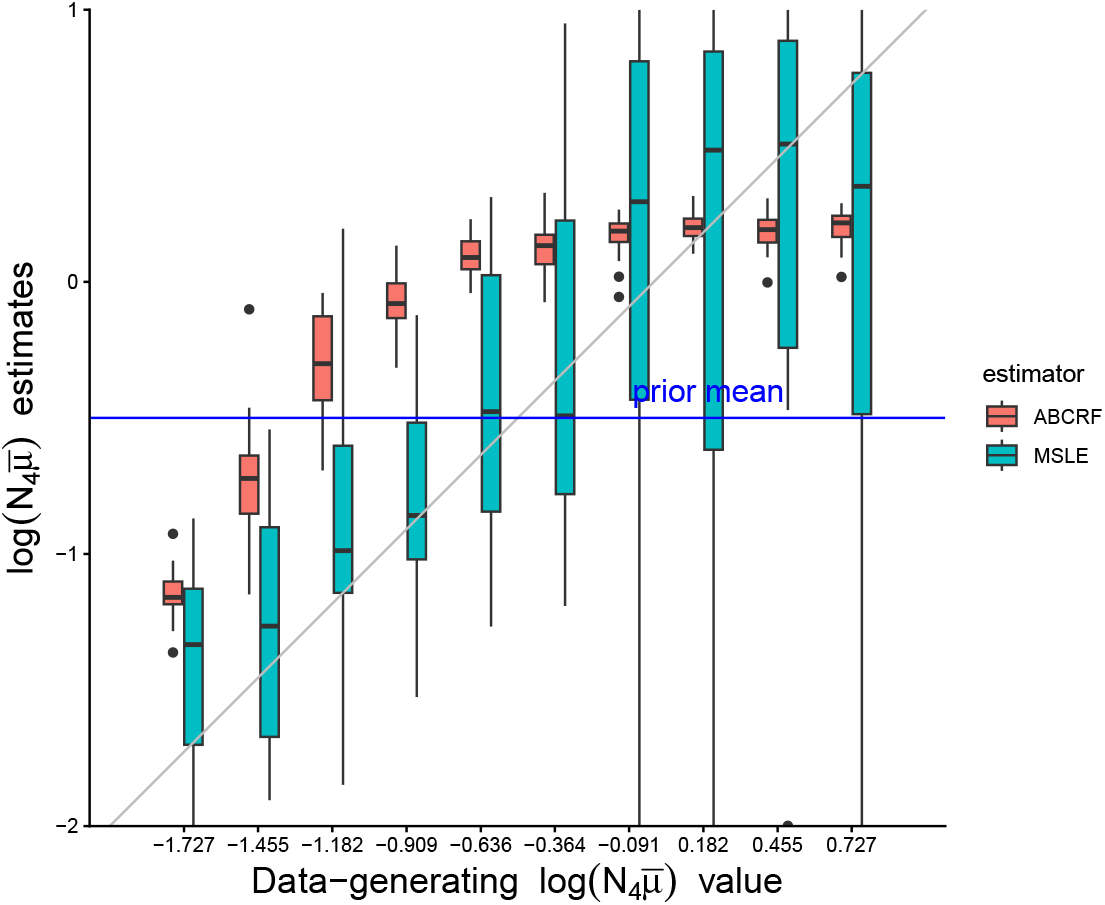
Distributions of MSLE and ABC-RF 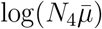 estimates for the third set of parameter values. Twenty simulated samples were analyzed for each given 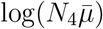 value. Other parameter values were constant as given by the third set of parameter values in Table S.5.

**Table S.9:** Performance summaries for the invasive ladybird scenario for the third set of parameter values and variable 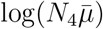. 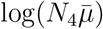 parameter values were varied as shown in Main Text Fig. S.3. 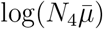 performance summaries are here average values over these variable 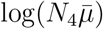 values. Other parameter values were constant as given by the third set of parameter values in Table S.5. Bias and root-mean-square error (RMSE) are reported for the maximum-summary likelihood estimates (MSLE), and the posterior mean (postEv) and posterior median (postMed) for ABC-RF. Coverage of confidence intervals with nominal 95% level is reported for summary-likelihood inference (profLR), for two forms of bootstrap correction described in the Text (bootLR and BcorCI), for the central intervals provided by ABC-RF (postCI), and for highest posterior density intervals (HPD CI). Bold font is used to emphasize for each parameter the bias value minimal in absolute value, the minimum RMSE, and the coverage closest to 0.95.

| parameter | Relative bias |  |  | Relative RMSE |  |  | coverage |  |  |  |  |
| --- | --- | --- | --- | --- | --- | --- | --- | --- | --- | --- | --- |
|  | MSLE | postEv | postMed | MSLE | postEv | postMed | profLR | bootLR | BcorCI | postCI | HPD CI |
| $\log(N_1\bar{\mu})$ | **-0.002** | 0.005 | 0.004 | 0.037 | 0.036 | **0.036** | **0.945** | 0.930 | 0.930 | 0.980 | 0.990 |
| $\log(N_2\bar{\mu})$ | **-0.013** | -0.023 | -0.031 | 0.055 | **0.044** | 0.051 | **0.935** | **0.935** | 0.930 | 0.985 | 0.990 |
| $\log(N_3\bar{\mu})$ | **-0.0005** | -0.115 | -0.097 | **0.087** | 0.128 | 0.112 | 0.975 | **0.960** | **0.960** | 1.000 | 1.000 |
| $\log(N_4\bar{\mu})$ | **0.013** | 0.122 | 0.135 | 0.241 | **0.200** | 0.208 | 1.000 | **0.970** | 0.990 | 1.000 | 0.990 |
| $t_1$ | **0.102** | 0.325 | 0.305 | **0.265** | 0.347 | 0.346 | 0.995 | 0.965 | 0.975 | **0.955** | 0.935 |
| $r_a$ | **0.020** | 0.091 | 0.061 | **0.095** | 0.132 | 0.116 | 0.960 | 0.940 | **0.945** | 0.985 | 0.985 |
| $\log(\bar{\mu})$ | **-0.010** | -0.036 | -0.031 | **0.050** | 0.060 | 0.058 | 0.960 | 0.940 | **0.945** | 1.000 | 1.000 |
| $\bar{p}$ | **0.037** | 0.048 | 0.040 | 0.171 | **0.157** | 0.162 | 0.960 | 0.945 | 0.935 | 0.965 | **0.950** |

##### S.4.2.5 Alternative parameter sampling rule

**Table S.10:** Performance summaries with alternative parameter sampling rule, for the invasive ladybird scenario. These results are based on running the inference workflow on the same 200 datasets as in Table S.7, simulated under the second set of parameters. Bias and root-mean-square error (RMSE) are reported for the maximum-summary likelihood estimates (MSLE), and the posterior mean (postEv) and posterior median (postMed) for ABC-RF. Coverage of confidence intervals with nominal 95% level is reported for summary-likelihood inference (profLR), for two forms of bootstrap correction described in the Text (bootLR and BcorCI), for the central intervals provided by ABC-RF (postCI), and for highest posterior density intervals (HPD CI). Bold font is used to emphasize for each parameter the bias value minimal in absolute value, the minimum RMSE, and the coverage closest to 0.95.

| parameter | Relative bias |  |  | Relative RMSE |  |  | coverage |  |  |  |  |
| --- | --- | --- | --- | --- | --- | --- | --- | --- | --- | --- | --- |
|  | MSLE | postEv | postMed | MSLE | postEv | postMed | profLR | bootLR | BcorCI | postCI | HPD CI |
| $\log(N_1\bar{\mu})$ | **0.007** | 0.012 | 0.011 | 0.037 | 0.037 | **0.037** | 0.975 | **0.955** | **0.955** | 0.940 | 0.990 |
| $\log(N_2\bar{\mu})$ | **-0.006** | -0.069 | -0.063 | **0.050** | 0.085 | 0.086 | **0.955** | 0.925 | 0.925 | 1.000 | 1.000 |
| $\log(N_3\bar{\mu})$ | **0.020** | -0.050 | -0.030 | 0.099 | 0.087 | **0.086** | 0.975 | **0.955** | **0.955** | 1.000 | 1.000 |
| $\log(N_4\bar{\mu})$ | **-0.011** | -0.036 | -0.045 | 0.139 | **0.071** | 0.078 | 0.980 | **0.955** | 0.960 | 1.000 | 1.000 |
| $t_1$ | **0.066** | 0.150 | 0.157 | 0.303 | **0.160** | 0.172 | **0.960** | 0.920 | **0.940** | 1.000 | 1.000 |
| $r_a$ | **0.094** | 0.102 | 0.099 | 0.250 | **0.146** | 0.156 | 0.985 | **0.975** | **0.975** | 1.000 | 0.985 |
| $\log(\bar{\mu})$ | **-0.006** | -0.066 | -0.055 | **0.051** | 0.087 | 0.082 | 0.970 | **0.945** | 0.935 | 1.000 | 1.000 |
| $\bar{p}$ | **0.011** | 0.026 | 0.013 | 0.156 | **0.142** | 0.143 | 0.975 | 0.970 | **0.965** | 0.980 | **0.965** |

### S.5 Human admixture scenario

#### S.5.1 Details of simulation scenario

##### S.5.1.1 The demo-genetic model

The Human admixture scenarios considered in our study are sub-models of a scenario with 17 variable parameters studied by Raynal *et al*. (2019) and Collin *et al*. (2021, scenario 2) to illustrate the ABC-RF method. This scenario includes (i) an ancient demographic change in the ancestral African population, (ii) a single out-of-Africa colonization event giving an ancestral out-of-Africa population which secondarily split into one European and one East Asian population, and (iii) a recent genetic admixture of the Americans of African Ancestry in SW-USA between their African ancestors and individuals of European origin. The original 17-parameter scenario is detailed in Figure 1 (right panel) of the Main Text. In this highly parameterized scenario, very few original parameters can be estimated. For this reason, we have chosen to derive sub-models in which we have fixed the number of variable parameters to 7 or 13, by fixing 10 or 4 parameters respectively, as detailed below.

Sample simulations for this scenario have been run using `diyabc` as in Raynal *et al*. (2019) and as described for the previous scenario. Notable differences compared to Raynal *et al*. (2019) lie in the way by which the event times have been expressed and in the way the `diyabc` software is used. In its basic use, the `diyabc` software draws values from parameter prior distributions until they satisfy a given set of constraints on the parameters (such as *t*_1_ *< t*_2_ *< t*_3_ *< t*_4_ in our scenario). A non-standard use of the `diyabc` simulator (based on the `-o` option in the online command of the executable file) is to provide it with a table of parameter values that do not depend on its rejection sampling and are therefore fully controlled by the user, possibly with different rules than the rejection sampling described above. Further, one can reparametrize the model according to differences between split or merge time events *t*_12_ = *t*_2_ − *t*_1_, *t*_23_ = *t*_3_ − *t*_2_, *t*_34_ = *t*_4_ − *t*_3_, and sample these new parameters only under the constraint that they are positive. This allows more transparent control of the distribution of sampled parameter points by the inference workflow. Hence this reparametrization was used for SL inferences.

In the parametrization of the 13-parameter Human admixture scenario by bottleneck parameters, *b*_3_ = *d*_3_*/N*_bn3_, *b*_4_ = *d*_4_*/N*_bn4_, and *b*_34_ = *d*_34_*/N*_bn34_, their possible values are subject to further logical constraints with respect to the times of events in the demographic scenario, which can be represented as follows

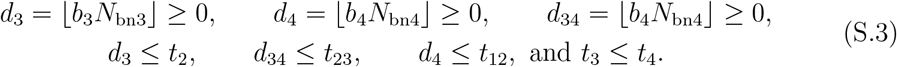

The choice of either sampling rules (rejection versus re-parametrization) has different implications for the two inference methods: rejection sampling under the constraint that *t*_1_ *< t*_2_ *< t*_3_ *< t*_4_ means that the sampled prior distributions of parameters are not uniform and have relatively low density near the boundaries defined by the non-box constraints (e.g., near *t*_2_ = *t*_3_). This tends to push posterior-based point estimates away from the boundary (i.e. when using ABC-RF), in comparison to posterior estimates based on uniform priors. Using the reparametrization instead of rejection sampling avoids this effect. The maximum-likelihood estimates obtained by our summary-likelihood method are comparatively insensitive to this effect of rejection sampling and can be much closer to the boundary. Hence, for easier comparison of SL and ABC-RF results, the reparametrization was also used for building the reference tables used in ABC-RF inferences.

The SNP data associated to those human admixture scenarios were obtained from individuals originating from four Human populations (30 unrelated individuals per population) using the freely accessible public 1000 Genome databases (i.e. the VCF format files including variant calls available at http://www.1000genomes.org/data; The 1000 Genomes Project Consortium, 2012). The four Human populations included the Yoruba population (Nigeria), encoded YRI in the 1000 genome database; the Han Chinese population (China), encoded CHB; the British population (England and Scotland), encoded GBR; and the population composed of Americans of African Ancestry in SW-USA, encoded ASW. Five or ten thousands independent SNP loci were selected from the 22 autosomal chromosomes using the criteria described in the Appendix S2 of Collin *et al*. (2021). In particular, only SNPs with a minimum allele frequency ≥ 0.01 were retained, and the same constraint was applied in the `diyabc` simulations. In the reference tables used for our inferences, the raw SNP genotype data for the scenarios studied are summarized by a total of 130 statistical summaries described in section 2.6.3 of the `diyabc` documentation (Collin *et al*., 2021; https://diyabc.github.io/doc/).

##### S.5.1.2 Data-generating parameter values

For the 13-parameter variant of the Human admixture scenario with composite bottleneck parameters, the data-generating values for the estimated parameters are log(*N*_1_) = 4.7, log(*N*_2_) = 3.4, *t*_1_ = 6, *r*_a_ = 0.21, log(*t*_12_) = 2.2, *d*_3_*/*Nbn_3_ = 0.26, log(Nbn_3_) = 2.1, *d*_4_*/*Nbn_4_ = 0.1, log(Nbn_4_) = 2.3, log(1 + *t*_23_) = 1.3, *d*_34_*/*Nbn_34_ = 0.44, log(Nbn_34_) = 1.5, log(*N*_a_) = 2.7, and those for known, non-estimated parameters are *N*_3_ = 6310, *N*_4_ = 3162 *N*_34_ = 1259 and *t*_4_ = 1007.

For the other 13-parameter variant, without composite parameters, the data-generating values are log(*N*_1_) = 3.2, log(*N*_2_) = 3.5, log(*N*_3_) = 3.8, log(*N*_4_) = 3.5, *t*_1_ = 6, *r*_a_ = 0.17, log(*t*_12_) = 2.2, *d*_3_ = 42, *d*_4_ = 9, log(1 + *t*_23_) = 1.7, *d*_34_ = 24, log(1 + *t*_34_) = 2.9, log(*N*_a_) = 2.7 for estimated ones; and *N*_bn3_ = 160, *N*_bn4_ = 98, *N*_34_ = 1259, and *N*_bn34_ = 74 for fixed known ones.

For the 7-parameter variant of the Human admixture scenario, the data-generating values are log(*N*_2_) = 3.5, *t*_1_ = 6, log(*t*_12_) = 2.2, log(1 + *t*_23_) = 1.5, Nbn_34_ = 65, log(1 + *t*_34_) = 3, log(*N*_a_) = 2.7 for estimated ones, and *N*_1_ = 12589, *N*_3_ = 19953, *N*_4_ = 3162, *r*_a_ = 0.2, *d*_3_ = 42, Nbn_3_ = 160, *d*_4_ = 9, Nbn_4_ = 98, *N*_34_ = 1259, *d*_34_ = 24 for fixed known ones.

##### S.5.1.3 Sampled parameter ranges for Human admixture scenarios

In the 7-parameters scenario, the ranges of sampled parameter space were set to [3, 5] for log(*N*_2_), [1, 30] for *t*_1_, [log(50), log(5000)] for log(*t*_12_), [0, log(5001)] for log(1 + *t*_23_) and log(1 + *t*_34_), [5, 500] for *N*_bn34_, and [2, 4] for log(*N*_a_).

In the 13-parameters scenario without composite bottleneck parameters, additional ranges were set to [3, 5] for all log(*N*_*i*_), [1, 50] for *d*_3_ and *d*_4_, and [0, 50] for *d*_34_.

In the 13-parameters scenario with composite bottleneck parameters, the ranges were [3, 5] for log(*N*_1_) and log(*N*_2_), [1, 30] for *t*_1_, [0.05, 0.95] for *r*_a_, [log(50), log(900)] for log(*t*_12_), [0, 1] for the three composite bottleneck parameters, [log(50), log(5000)] for log(*N*_bn3_), log(*N*_bn4_), and log(*N*_bn34_), [0, log(901)] for log(1 + *t*_23_), and [2, 4] for log(*N*_a_).

#### S.5.2 7-parameter inferences: additional simulation results

##### S.5.2.1 Distributions of p-values

**Figure S.4:**
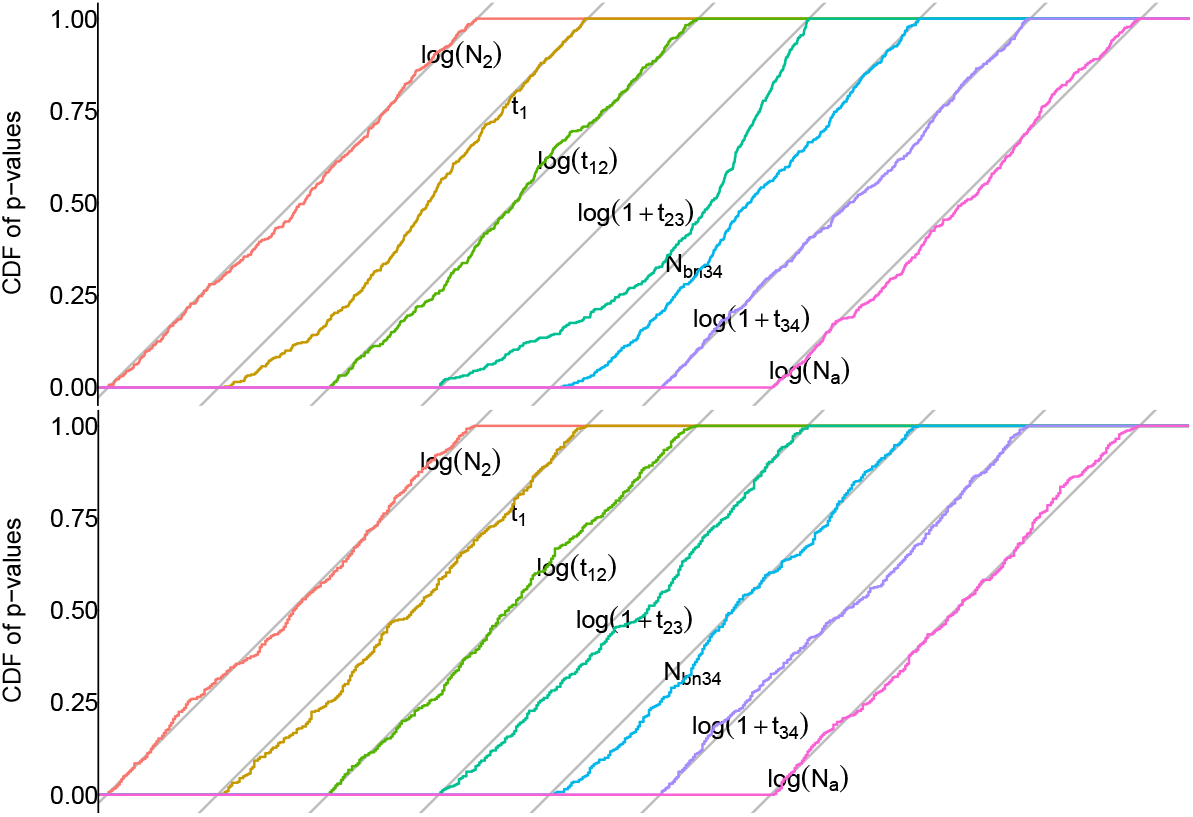
Cumulative distributions of p-values of summary-LRTs for the 7-parameter human admixture scenario. This Figure describes the same simulations as in Main Text Table 3. The distributions of each uncorrected (top) and bootstrap-corrected (bottom) tests for each parameter is shown relative to its own 1:1 reference line.

##### S.5.2.2 Comparison with analyses of datasets of 10000 SNPs

**Table S.11:**
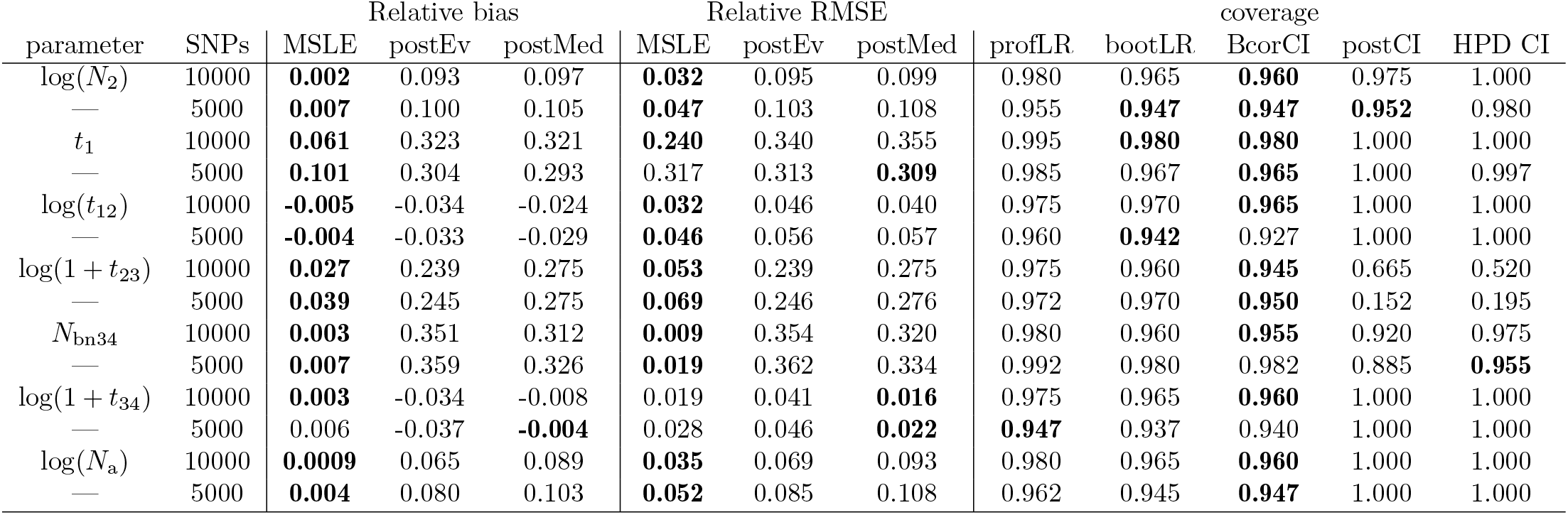
Performance summaries for the 7-parameter Human admixture scenario. Alternating rows show results for datasets of 10000 SNPs (200 simulated datasets) compared to results from Table S.4 for datasets of 5000 SNP (400 simulated datasets). Bias and root-mean-square error (RMSE) are reported for the maximum-summary likelihood estimates (MSLE), and the posterior mean (postEv) and posterior median (postMed) for ABC-RF. Coverage of confidence intervals with nominal 95% level is reported for summary-likelihood inference (profLR), for two forms of bootstrap correction described in the Text (bootLR and BcorCI), for the central intervals provided by ABC-RF (postCI), and for highest posterior density intervals (HPD CI). Bold font is used to emphasize for each parameter the bias value minimal in absolute value, the minimum RMSE, and the coverage closest to 0.95.

##### S.5.2.3 Boundary effects

Table S.12 reports results of inferences for two series of 400 datasets simulated as in Table 3 except that *N*_bn34_ = 5 in the first series (top seven rows of performance results) and 500 in the second series (bottom seven rows). The lines for the *N*_bn34_ shows the distinct coverage of the two credibility intervals compared.

**Table S.12:**
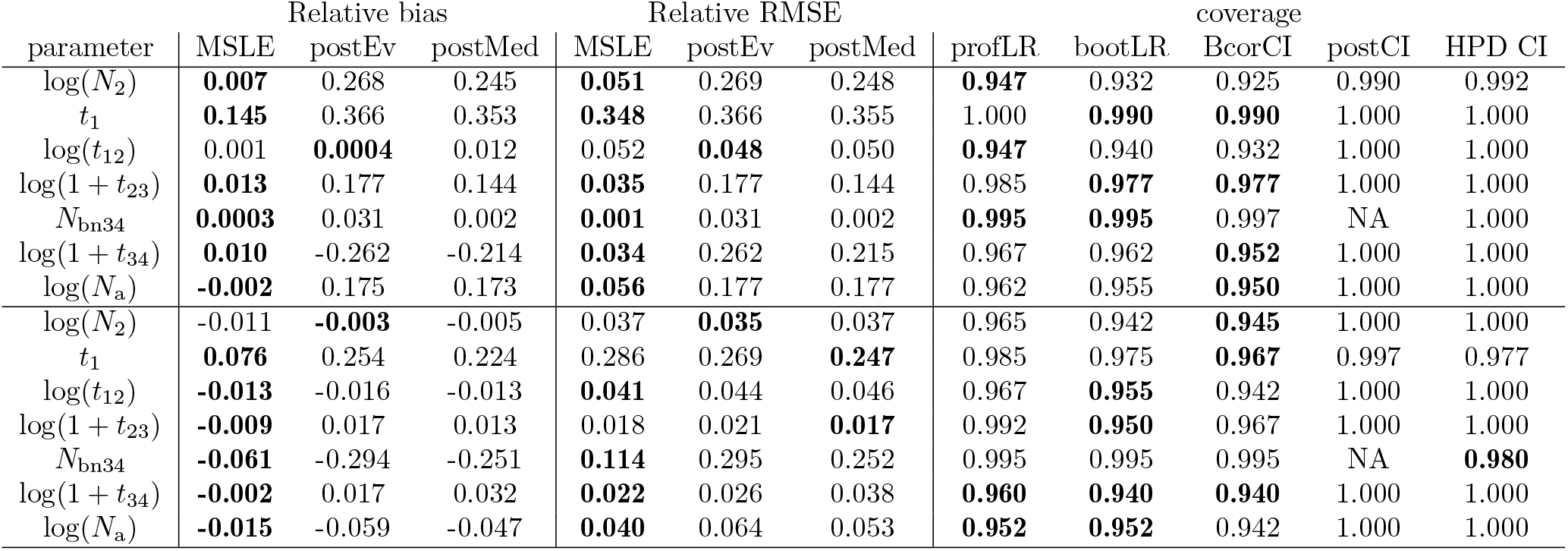
Performance summaries with *N*_bn34_ at either bound of its range. Bias and root-mean-square error (RMSE) are reported for the maximum-summary likelihood estimates (MSLE), and the posterior mean (postEv) and posterior median (postMed) for ABC-RF. Coverage of confidence intervals with nominal 95% level is reported for summary-likelihood inference (profLR), for two forms of bootstrap correction described in the Text (bootLR and BcorCI), for the central intervals provided by ABC-RF (postCI), and for highest posterior density intervals (HPD CI). Bold font is used to emphasize for each parameter the bias value minimal in absolute value, the minimum RMSE, and the coverage closest to 0.95.

| parameter | Relative bias |  |  | Relative RMSE |  |  | coverage |  |  |  |  |
| --- | --- | --- | --- | --- | --- | --- | --- | --- | --- | --- | --- |
|  | MSLE | postEv | postMed | MSLE | postEv | postMed | profLR | bootLR | BcorCI | postCI | HPD CI |
| $\log(N_2)$ | **0.007** | 0.268 | 0.245 | **0.051** | 0.269 | 0.248 | **0.947** | 0.932 | 0.925 | 0.990 | 0.992 |
| $t_1$ | **0.145** | 0.366 | 0.353 | **0.348** | 0.366 | 0.355 | 1.000 | **0.990** | **0.990** | 1.000 | 1.000 |
| $\log(t_{12})$ | 0.001 | **0.0004** | 0.012 | 0.052 | **0.048** | 0.050 | **0.947** | 0.940 | 0.932 | 1.000 | 1.000 |
| $\log(1 + t_{23})$ | **0.013** | 0.177 | 0.144 | **0.035** | 0.177 | 0.144 | 0.985 | **0.977** | **0.977** | 1.000 | 1.000 |
| $N_{\text{bn}34}$ | **0.0003** | 0.031 | 0.002 | **0.001** | 0.031 | 0.002 | **0.995** | **0.995** | 0.997 | NA | 1.000 |
| $\log(1 + t_{34})$ | **0.010** | -0.262 | -0.214 | **0.034** | 0.262 | 0.215 | 0.967 | 0.962 | **0.952** | 1.000 | 1.000 |
| $\log(N_a)$ | **-0.002** | 0.175 | 0.173 | **0.056** | 0.177 | 0.177 | 0.962 | 0.955 | **0.950** | 1.000 | 1.000 |
| $\log(N_2)$ | -0.011 | **-0.003** | -0.005 | 0.037 | **0.035** | 0.037 | 0.965 | 0.942 | **0.945** | 1.000 | 1.000 |
| $t_1$ | **0.076** | 0.254 | 0.224 | 0.286 | 0.269 | **0.247** | 0.985 | 0.975 | **0.967** | 0.997 | 0.977 |
| $\log(t_{12})$ | **-0.013** | -0.016 | -0.013 | **0.041** | 0.044 | 0.046 | 0.967 | **0.955** | 0.942 | 1.000 | 1.000 |
| $\log(1 + t_{23})$ | **-0.009** | 0.017 | 0.013 | 0.018 | 0.021 | **0.017** | 0.992 | **0.950** | 0.967 | 1.000 | 1.000 |
| $N_{\text{bn}34}$ | **-0.061** | -0.294 | -0.251 | **0.114** | 0.295 | 0.252 | 0.995 | 0.995 | 0.995 | NA | **0.980** |
| $\log(1 + t_{34})$ | **-0.002** | 0.017 | 0.032 | **0.022** | 0.026 | 0.038 | **0.960** | **0.940** | **0.940** | 1.000 | 1.000 |
| $\log(N_a)$ | **-0.015** | -0.059 | -0.047 | **0.040** | 0.064 | 0.053 | **0.952** | **0.952** | 0.942 | 1.000 | 1.000 |

##### S.5.2.4 Results with synthetic reference table

The fit of the simulated joint distribution for parameters and summary statistics in a reference table allows, in principle, the likelihood to be evaluated following eq. 3 for any dataset simulated under the same model as the reference table. Thus, a single such fit could in principle be used to perform inference for different real or simulated datasets 𝒟. However, in practice, attempts to reuse reference tables built for other datasets may not be successful, if the likelihood evaluation is precise only after several iterations where the sampling of new parameter points is driven by the analyzed dataset.

**Table S.13:** Performance of inferences using synthetic reference table. Bias and root-mean-square error (RMSE) are reported for the maximum-summary likelihood estimates (MSLE). Coverage of confidence intervals with nominal 95% level is reported for summary-likelihood inference (profLR), and for two forms of bootstrap correction described in the Text (bootLR and BcorCI). Bold font is used to emphasize for each parameter the coverage closest to 0.95.

| parameter | rel. bias | rel. RMSE | profLR | bootLR | BcorCI |
| --- | --- | --- | --- | --- | --- |
| $\log(N_2)$ | -0.036 | 0.114 | 0.840 | **0.850** | 0.835 |
| $t_1$ | 0.525 | 0.616 | **0.595** | 0.435 | 0.445 |
| $\log(t_{12})$ | -0.061 | 0.105 | 0.670 | **0.675** | 0.650 |
| $\log(1 + t_{23})$ | 0.051 | 0.113 | **0.880** | 0.850 | 0.815 |
| $N_{\text{bn}34}$ | -0.011 | 0.140 | **0.955** | 0.925 | 0.925 |
| $\log(1 + t_{34})$ | 0.017 | 0.102 | 0.725 | **0.770** | 0.760 |
| $\log(N_a)$ | 0.088 | 0.200 | 0.505 | **0.535** | 0.495 |

**Figure S.5:**
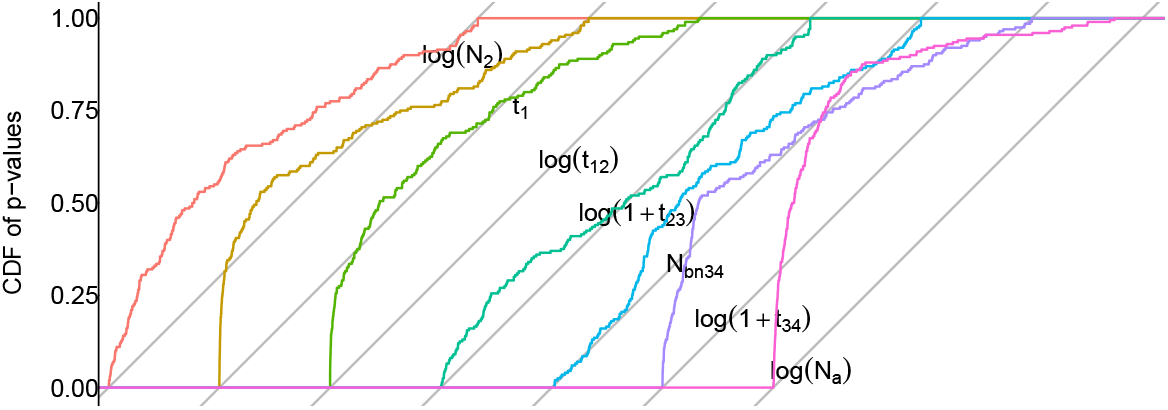
Distributions of p-values of summary-LRTs using a synthetic reference table.

In order to check whether this occurs, we built a single synthetic reference table of 20000 samples, by drawing 100 samples randomly from each of the 20000-sample reference tables (containing raw summary statistics) for each of the first 200 simulated datasets. This table was used to infer new projections and likelihood surfaces for the same 200 simulated datasets. Although the biases of resulting estimators are not necessarily large, the biases and RMSEs are uniformly larger than those of the original inferences, as can be seen in Table S.13 and Fig. S.5, and coverage is far from controlled. Thus, the inference workflow is best run essentially independently for each new dataset.

##### S.5.3.5 Comparison with simplified sampling rule

**Table S.14:** Performance using simplified sampling rule. These computations used the same 400 datasets as in Table 3 in the Main Text. Bias and root-mean-square error (RMSE) are reported for the maximum-summary likelihood estimates (MSLE). Coverage of confidence intervals with nominal 95% level is reported for summary-likelihood inference (profLR), and for two forms of bootstrap correction described in the Text (bootLR and BcorCI). Bold font is used to emphasize for each parameter the coverage closest to 0.95.

| parameter | rel. bias | rel. RMSE | coverage |  |  |
| --- | --- | --- | --- | --- | --- |
|  |  |  | profLR | bootLR | BcorCI |
| $\log(N_2)$ | 0.004 | 0.047 | **0.955** | **0.955** | 0.942 |
| $t_1$ | 0.114 | 0.329 | 0.970 | 0.962 | **0.957** |
| $\log(t_{12})$ | -0.006 | 0.046 | **0.945** | 0.937 | 0.930 |
| $\log(1 + t_{23})$ | 0.032 | 0.066 | 0.955 | 0.952 | **0.950** |
| $N_{\text{bn}34}$ | 0.006 | 0.019 | 0.992 | **0.977** | 0.987 |
| $\log(1 + t_{34})$ | 0.006 | 0.027 | **0.947** | 0.942 | 0.935 |
| $\log(N_a)$ | 0.001 | 0.052 | **0.950** | 0.942 | 0.940 |

**Table S.15:**
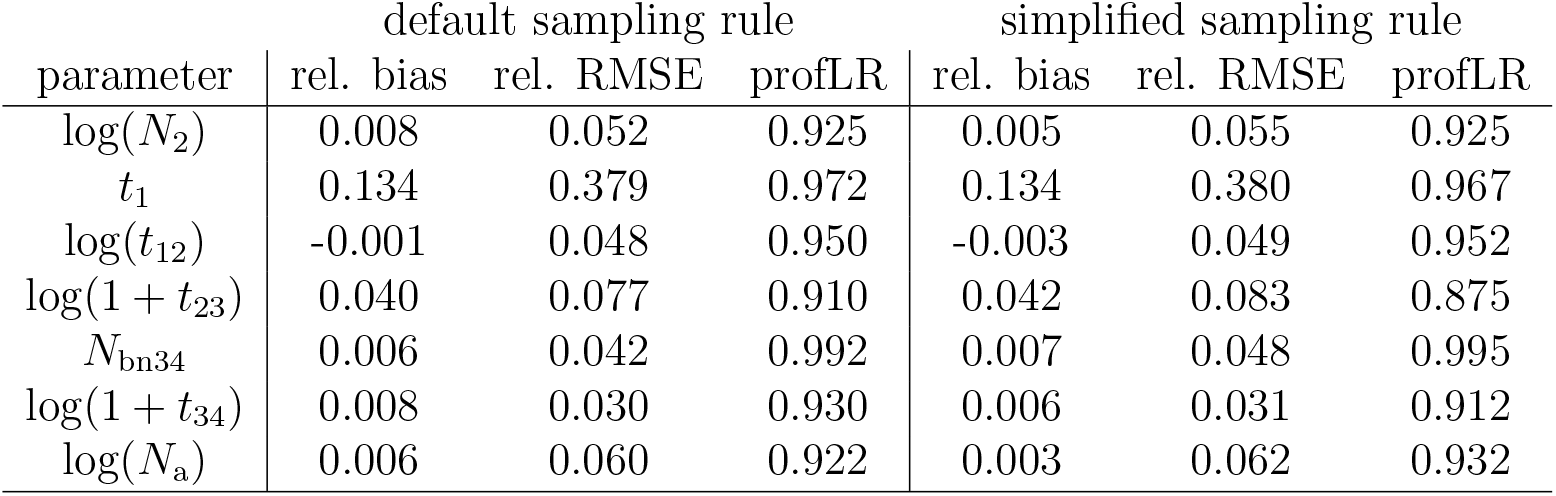
Performance of sampling rules on small reference tables. Bias, root-mean-square error (RMSE), and coverage of 95% intervals (profLR) are reported. All inferences are based on 6000-sample intermediate reference tables, built during the full 20000-sample inferences whose results are described in Tables 3 and S.14. There is only faint, if any, evidence of better performance when using the default sampling rule: marginally but consistently smaller RMSEs for all parameters; and a possibly faster convergence of coverage of profile summary-LRTs (“profLR”) for log(1 + *t*_23_), for which the importance of the iterative workflow is highlighted in the Main Text.

| parameter | default sampling rule |  |  | simplified sampling rule |  |  |
| --- | --- | --- | --- | --- | --- | --- |
|  | rel. bias | rel. RMSE | profLR | rel. bias | rel. RMSE | profLR |
| $\log(N_2)$ | 0.008 | 0.052 | 0.925 | 0.005 | 0.055 | 0.925 |
| $t_1$ | 0.134 | 0.379 | 0.972 | 0.134 | 0.380 | 0.967 |
| $\log(t_{12})$ | -0.001 | 0.048 | 0.950 | -0.003 | 0.049 | 0.952 |
| $\log(1 + t_{23})$ | 0.040 | 0.077 | 0.910 | 0.042 | 0.083 | 0.875 |
| $N_{\text{bn}34}$ | 0.006 | 0.042 | 0.992 | 0.007 | 0.048 | 0.995 |
| $\log(1 + t_{34})$ | 0.008 | 0.030 | 0.930 | 0.006 | 0.031 | 0.912 |
| $\log(N_a)$ | 0.006 | 0.060 | 0.922 | 0.003 | 0.062 | 0.932 |

#### S.5.3 13-parameter inferences: additional simulation results

##### S.5.3.1 Distributions of p-values

**Figure S.6:**
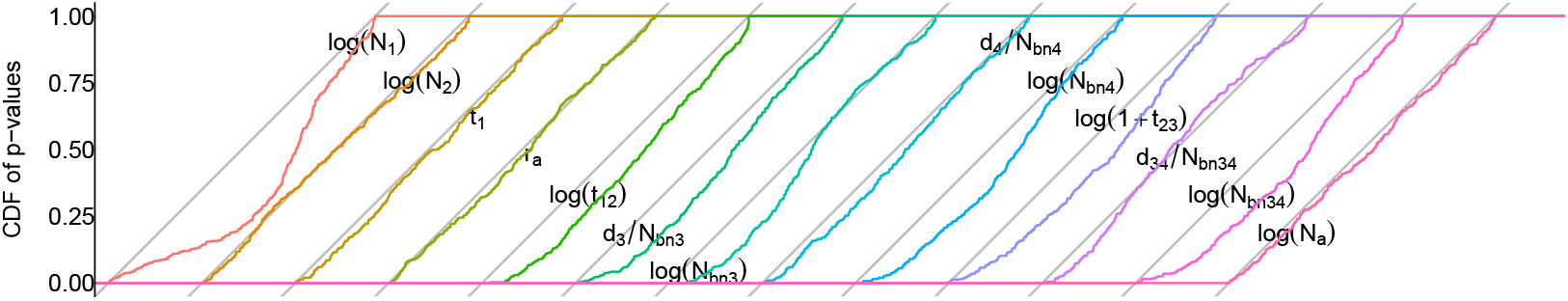
Distributions of p-values of summary-LRTs for the 13-parameter human admixture scenario. This Figure describes the same simulation results as in Table 4.

##### S.5.3.2 Interpretation of the compared RMSEs of summary-MLE and ABC-RF estimates

The RMSEs of summary ML estimates is lower or higher than that of ABC-RF estimates, depending on the parameter considered (Main Text Table 4). As emphasized in the Main Text, this pattern is expected and is affected by the position of data-generating values ***θ***^*†*^ relative to prior means, so such comparisons of RMSEs are not sufficient to characterize the relative estimation performance of the two methods. The RMSEs of ABC-RF estimators are also possibly affected by other effects illustrated in the main Text. Although we did not try to tell apart these different effects for all parameters, we can look for the distinctive pattern observed in the previous simulations, and interpreted as resulting for poor exploration of parameter space by the non-iterative workflow. This pattern, where for a given parameter the ABC-RF estimates are out of the range defined by the data-generating value and the prior mean, and have a variance much lower than that of summary-MLEs, is again observed for log(1 + *t*_23_), but also for log(*t*_12_), *N*_bn3_, log(*N*_bn4_), *b*_34_, all with substantially lower variance than the MSLE [all ratios of variances *<* 0.28 except 0.43 for log(*t*_12_)]. Each of these ABC-RF estimators is weakly correlated with the MSLE one for the same parameter [correlation *<* 0.5 except 0.57 for log(*t*_12_)].

Although log(*N*_a_) appears relatively well estimated by both methods, its ABC-RF estimator also meets these diagnostic criteria, except that it has higher correlation with the MSLE (0.75).

The admixture rate *r*_a_, population size *N*_2_, and the composite parameters *b*_3_ and *b*_4_ appear well inferred by both methods. However, the ABC-RF estimator of log(*N*_2_) has a suspiciously low variance ratio (0.33, versus between 0.61 and 1.31 for the three other parameters), suggesting either a strong effect of the prior mean on estimates, or an artifact of poor exploration of parameter space. The deficiency of low p-values for *b*_3_ may result from the constraint *d*_3_ ≥ 0 on the parameter space and the lack of information about *N*_bn3_ on this boundary (See Section S.5.3.5 below for a detailed analysis).

*t*_1_, *N*_1_ and *N*_bn34_ are poorly estimated by both methods. This is less immediately obvious for *N*_1_ and *N*_bn34_, as their ABC-RF estimators have low RMSE, but in these cases the data-generating value *θ*^*†*^ is close to the prior mean. Fig. S.7 shows that the point estimates for both log(*N*_1_) and log(*N*_bn34_) remain close to their prior mean in the reference table, independently of the data-generating value, a result expected when the data provide not information about the parameter. This also explains why the posterior estimates have much lower variance than summary-MLEs (variance ratios 0.025 and 0.039). For log(*N*_bn34_), this interpretation may be overlooked as the prior mean in the reference table differs from the mid-range because of the additional constraints on parameters (Main Text eq. S.3).

**Figure S.7:**
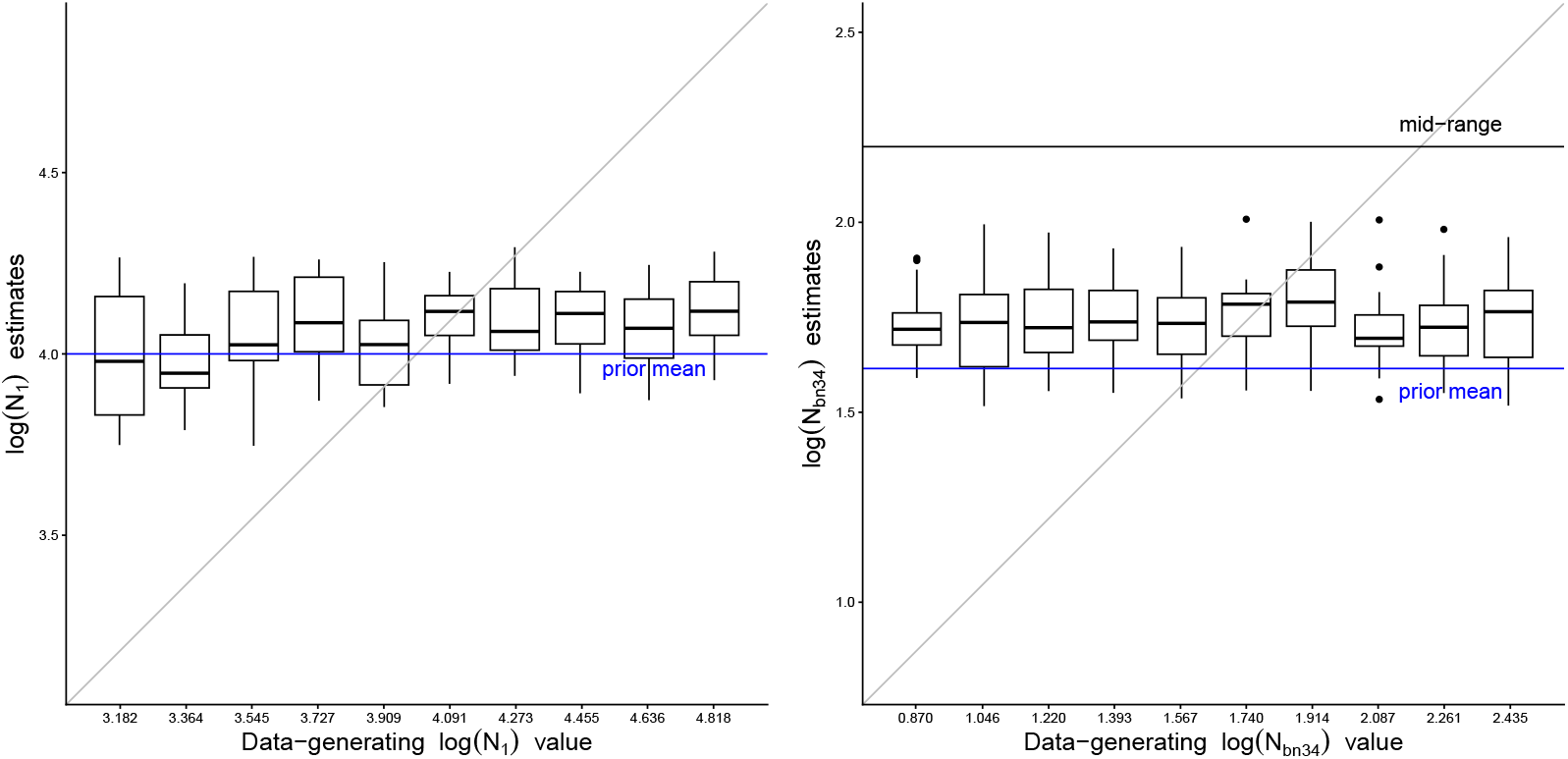
Distributions of ABC-RF estimates for the parameters log(*N*_1_) (left) and log(*N*_bn34_) (right). Twenty simulated samples were analyzed for each given value of the variable parameter. Other parameters are set as in Table 4, except that in the simulations for variable *N*_bn34_, the data-generating *t*_23_ value was increased (log(1 + *t*_23_) = 2.5) in order to increase the range of log(*N*_bn34_) values compatible with the parameter constraints for this model (Main Text eq. S.3).

##### S.5.3.3 Real-data inferences

**Table S.16:**
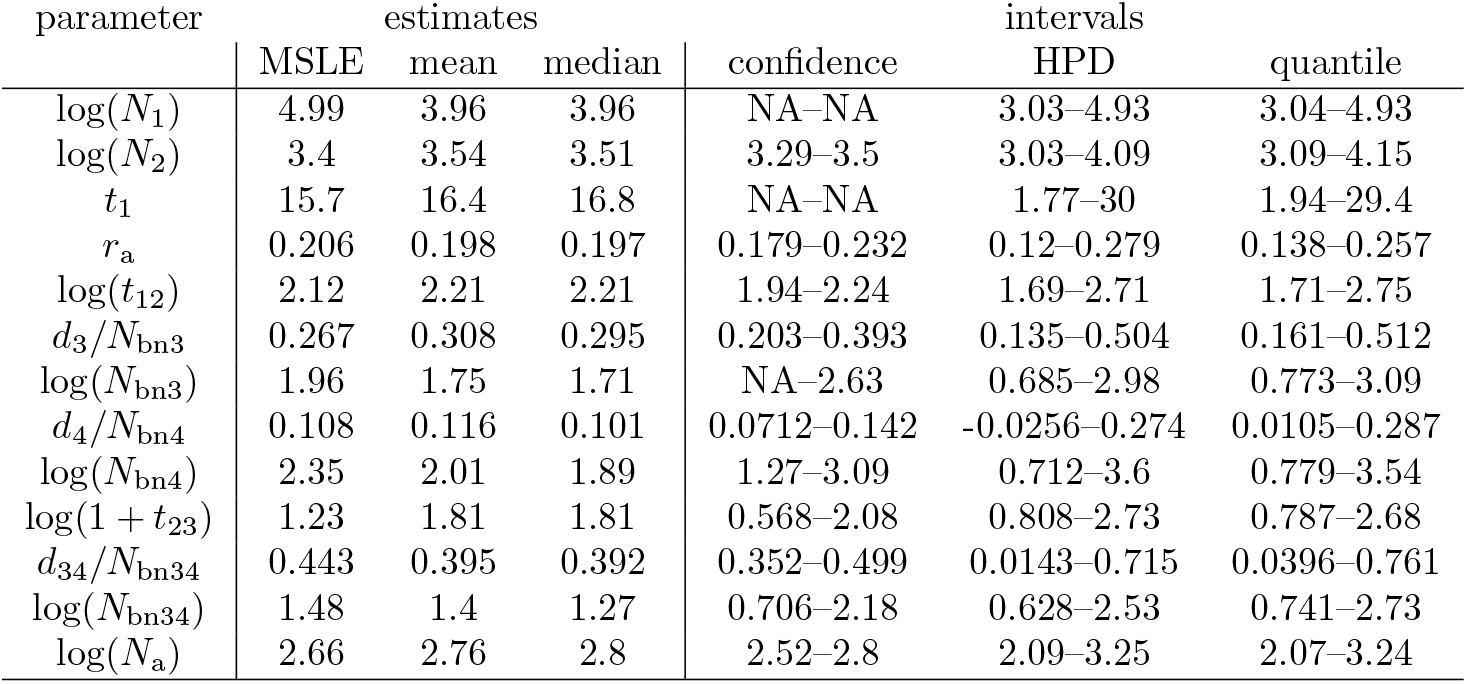
Inferences from real data. 5,000 SNPs were used. The inferences were conducted exactly as for the simulated data for the 13-parameter base case (the one with composite bottleneck parameters). A “NA” lower or upper bound of a confidence interval means that the lower or upper bound of the explored parameter range was not excluded from the interval.

##### S.5.3.4 Reference tables of 50,000 samples

In general, poor estimation of some parameters can be interpreted as demonstrating that there is weak information about them in the data, or that the estimation method fails to capture such information. To test the possibility that not enough iterations of the summary-likelihood workflow have been run to fill the top region of the likelihood surface, we increased the size of the reference tables from 38000 to 50000 simulated samples (i.e., 24 additional iterations), for the first 200 simulated datasets. The effect of these additional iterations is, if any, too small to be detectable (Table S.17).

**Table S.17:** Performance of summary-likelihood for the 13-parameter Human admixture scenario. To obtain these results, the inferences initially performed on the first 200 of the 400 datasets considered in Table 4, were run for more iterations, extending the reference tables to 50,000 simulations. Bias and root-mean-square error (RMSE) are reported for the maximum-summary likelihood estimates (MSLE). Coverage of confidence intervals with nominal 95% level is reported for summary-likelihood inference (profLR), and for two forms of bootstrap correction described in the Text (bootLR and BcorCI). Bold font is used to emphasize for each parameter the coverage closest to 0.95. There is no detectable overall effect of extending the reference table on the bias and RMSE.

| parameter | rel. bias | rel. RMSE | profLR | bootLR | BcorCI |
| --- | --- | --- | --- | --- | --- |
| $\log(N_1)$ | -0.118 | 0.369 | 0.975 | **0.950** | **0.950** |
| $\log(N_2)$ | -0.007 | 0.033 | 0.970 | **0.945** | **0.945** |
| $t_1$ | 0.045 | 0.229 | 0.985 | 0.925 | **0.950** |
| $r_a$ | -0.001 | 0.014 | 0.980 | 0.940 | **0.945** |
| $\log(t_{12})$ | -0.021 | 0.056 | 1.000 | 0.970 | **0.965** |
| $d_3/N_{\text{bn}3}$ | 0.006 | 0.032 | 1.000 | **0.995** | **0.995** |
| $\log(N_{\text{bn}3})$ | -0.076 | 0.155 | 0.995 | **0.980** | 0.990 |
| $d_4/N_{\text{bn}4}$ | 0.001 | 0.022 | 0.975 | **0.945** | **0.955** |
| $\log(N_{\text{bn}4})$ | -0.016 | 0.191 | 0.980 | 0.935 | **0.950** |
| $\log(1 + t_{23})$ | 0.024 | 0.114 | 0.980 | **0.960** | 0.975 |
| $d_{34}/N_{\text{bn}34}$ | -0.033 | 0.055 | 0.995 | **0.985** | 0.995 |
| $\log(N_{\text{bn}34})$ | -0.001 | 0.133 | 0.985 | **0.960** | **0.960** |
| $\log(N_a)$ | -0.012 | 0.041 | 0.960 | **0.955** | 0.960 |

##### S.5.3.5 Analyzing the performance of summary-LRTs for the composite parameter *d*_3_*/N*_bn3_

In Main Text Table 4, *d*_3_*/N*_bn3_ appears estimable with some precision. Yet the confidence intervals appear to have higher than nominal coverage, which suggests that there is little information about this composite bottleneck intensity parameter. These observations can be reconciled as follows. Short bottlenecks have little impact on genetic polymorphism. Thus, for the lowest values of bottleneck length *d*_3_, there is little power to distinguish strong versus weak bottlenecks (corresponding to low versus high *N*_bn3_). For *d*_3_ = 0 in particular the log-likelihood should be a constant *ℓ*_0_, independent of *N*_bn3_. Further, for datasets whose summary-MLE for *d*_3_ approaches the boundary, the profile likelihood for *d*_3_ = 0 (or *d*_3_ = 1) may not be much lower than the likelihood maximum (for simplicity of exposition, we consider the likelihood only a function of *d*_3_ and *N*_bn3_ here). These patterns are illustrated in Fig. S.8. Since the likelihood isoline for the tested value 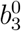 of *b*_3_ = *d*_3_*/N*_bn3_ has a point at the boundary, the profile likelihood for 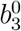 is at least the log-likelihood *ℓ*_0_ at the boundary. Hence, the likelihood ratio statistic for the value 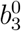 of the composite bottleneck parameter should be small, and the power of the test should be low, when the summary-MLE for bottleneck length *d*_3_ approaches the boundary.

To check this interpretation, we performed additional simulations in which we reduced the number of estimated parameters but increased the number of simulated datasets (500, including the 200 of the original 13-parameter inferences). Only *t*_12_, *N*_a_ and the two bottleneck parameters *d*_3_*/N*_bn3_ and log(*N*_bn3_) were estimated, the other ones being fixed to their true values. The deficiency of low p-values is retained in these simulations (Table S.18).

**Table S.18:** Performance of summary-likelihood for 4-parameters inferences. Details of simulations are as for the 13-parameter inferences, except that the log(*N*_bn3_) range was extended to [0, log(50000)], and the inference for each dataset is based on final references tables of 11000 samples. Bias and root-mean-square error (RMSE) are reported for the maximum-summary likelihood estimates (MSLE). Coverage of confidence intervals with nominal 95% level is reported for summary-likelihood inference (profLR), and for two forms of bootstrap correction described in the Text (bootLR and BcorCI). Bold font is used to emphasize for each parameter the coverage closest to 0.95.

| parameter | rel. bias | rel. RMSE | profLR | bootLR | BcorCI |
| --- | --- | --- | --- | --- | --- |
| $\log(t_{12})$ | 0.002 | 0.020 | 0.966 | 0.960 | **0.958** |
| $d_3/N_{\text{bn}3}$ | 0.008 | 0.053 | 0.996 | **0.984** | 0.990 |
| $\log(N_{\text{bn}3})$ | -0.075 | 0.192 | **0.964** | 0.914 | 0.918 |
| $\log(N_a)$ | -0.0005 | 0.015 | 0.930 | **0.932** | 0.928 |

Since deficiencies of low p-values of the profile summary-LRT of 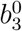 are expected more for datasets whose *d*_3_ estimates are low, we show separately in Fig. S.9 the distributions of p-values for datasets with low versus datasets with high *d*_3_ estimates, the threshold *d*_3_ being the data-generating value *d*_3_ = 32 in these simulations. There are deficiencies in both cases, but they are, as expected, stronger for low 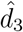 . For high 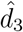, the bootstrap test approximately corrects the distribution.

**Figure S.8:**
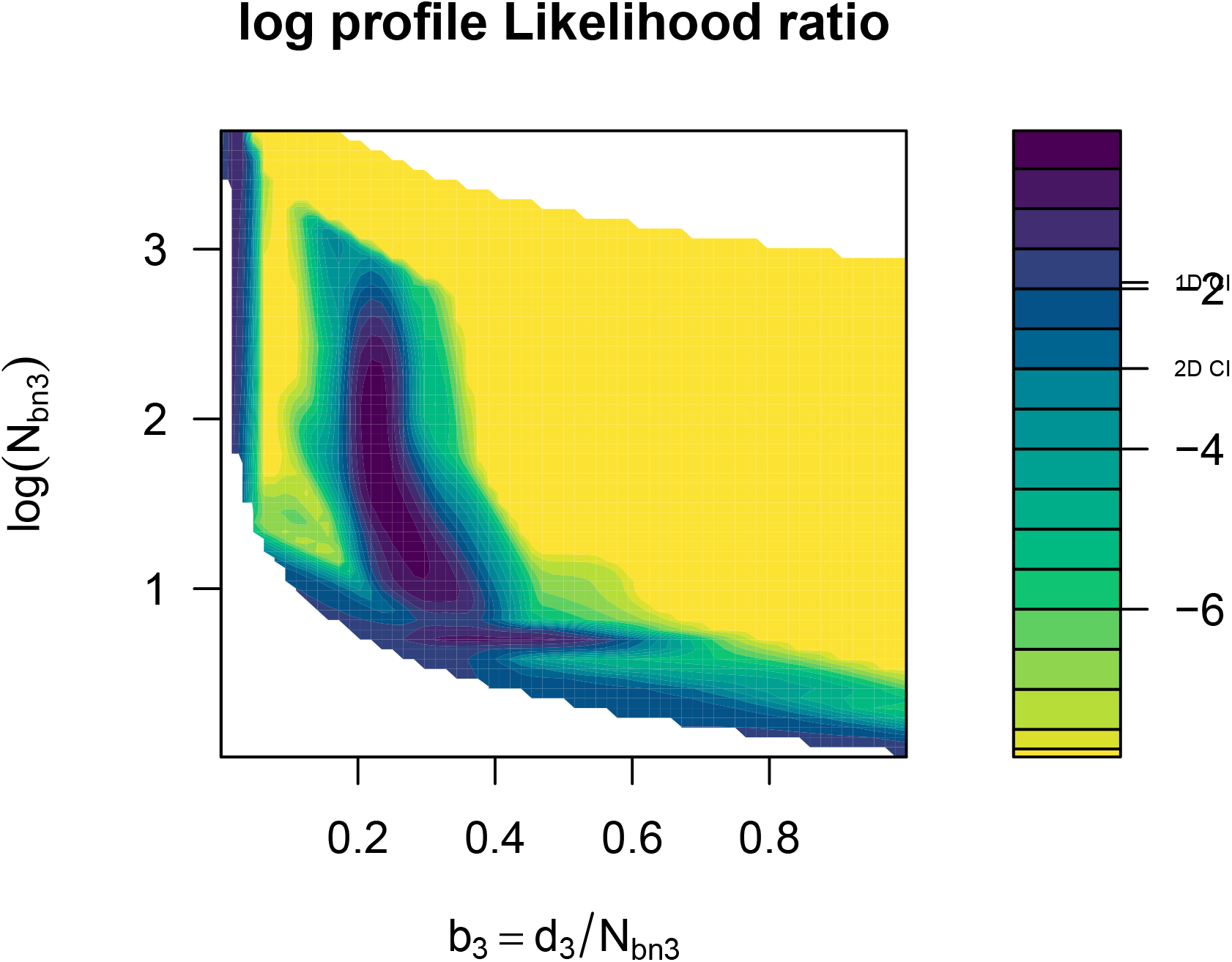
Illustrative likelihood surface profile for bottleneck parameters. This Figure illustrates a recurrent pattern for different simulated datasets: the narrow vertical ridge provides the 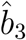 estimate near the true value 0.26, but likelihoods remain relatively high along the lower boundary of the plotted area, which corresponds to vanishing bottleneck length.

**Figure S.9:**
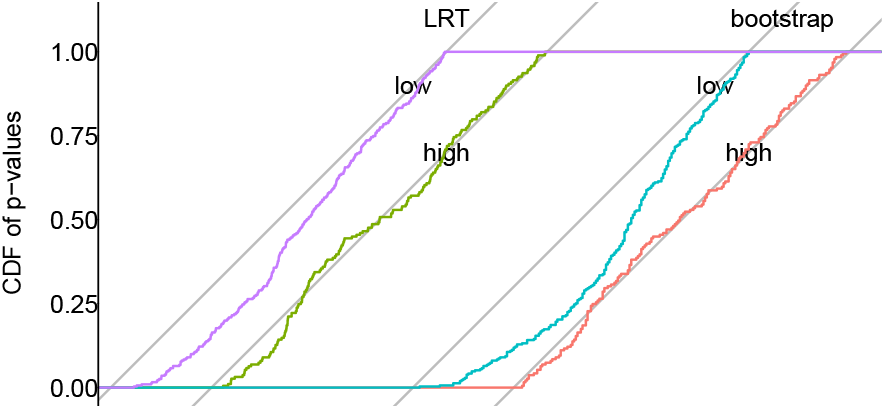
Distributions of p-values of tests for the composite parameter *d*_3_*/N*_bn3_. Distributions are shown for likelihood ratio (left) and bootstrap tests (right), distinguishing in each case the distributions for datasets with low estimates of bottleneck length, 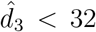, from those with higher estimates (189 and 311 datasets, respectively).

##### S.5.3.6 Inference without composite parameters

Results for the 13-parameter variant without composite bottleneck intensity parameters are detailed in Table S.19 and Fig. S.10. Only the admixture parameter *r*_a_ can be estimated precisely, with approximately uniform distribution of p-values. *d*_4_ comes second in terms of RMSE. At the opposite, MSLE estimators of *t*_1_, *t*_12_, and the population size parameters all have high relative RMSEs (from 0.195 to 0.528), and large deficiencies of low p-values. These results confirm the identifiability issues that led us to consider the alternative 13-parameter variant with composite bottleneck intensity parameters. By applying the same diagnostic criteria as in Section S.5.3.2, one can see that the ABC-RF results for log(*N*_1_), log(*N*_3_), log(1+*t*_23_) and log(1+*t*_23_) are suspect because the posterior-mean estimator falls on average out of the interval defined by the prior mean and the data-generating parameter value. All ABC-RF estimators except the one for *r*_a_ are suspect also in terms of their low variance relative to summary-MLEs: the variance ratios are 0.964 for *r*_a_, but 0.216 for *d*_4_, 0.14 for log(*N*_2_), 0.13 for *d*_34_, and *<* 0.06 for all other parameters.

**Figure S.10:**
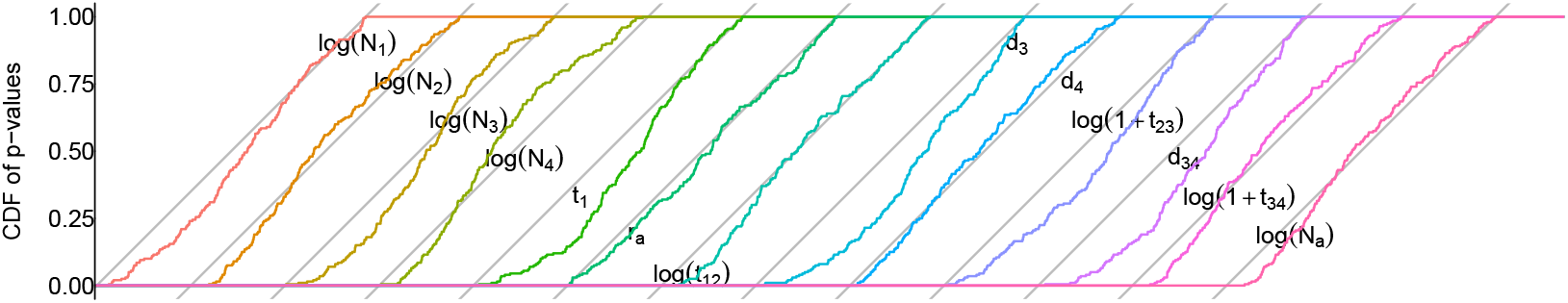
Distributions of p-values of summary-LRTs for the 13-parameter Human admixture scenario without composite parameters.

**Table S.19:** Performance for the 13-parameter Human admixture scenario without composite bottleneck parameters. Bias and root-mean-square error (RMSE) are reported for the maximum-summary likelihood estimates (MSLE), and the posterior mean (postEv) and posterior median (postMed) for ABC-RF. Coverage of confidence intervals with nominal 95% level is reported for summary-likelihood inference (profLR), for two forms of bootstrap correction described in the Text (bootLR and BcorCI), for the central intervals provided by ABC-RF (postCI), and for highest posterior density intervals (HPD CI). Bold font is used to emphasize for each parameter the bias value minimal in absolute value, the minimum RMSE, and the coverage closest to 0.95.

| parameter | Relative bias |  |  | Relative RMSE |  |  | coverage |  |  |  |  |
| --- | --- | --- | --- | --- | --- | --- | --- | --- | --- | --- | --- |
|  | MSLE | postEv | postMed | MSLE | postEv | postMed | profLR | bootLR | BcorCI | postCI | HPD CI |
| $\log(N_1)$ | **0.374** | 0.464 | 0.484 | 0.527 | **0.469** | 0.491 | 0.995 | 0.965 | 0.980 | **0.955** | 0.930 |
| $\log(N_2)$ | 0.151 | 0.141 | **0.113** | 0.200 | 0.150 | **0.124** | 1.000 | **0.990** | 0.995 | 1.000 | 1.000 |
| $\log(N_3)$ | 0.292 | 0.131 | **0.122** | 0.374 | **0.142** | 0.144 | 0.995 | **0.985** | 0.995 | 1.000 | 1.000 |
| $\log(N_4)$ | 0.325 | 0.253 | **0.248** | 0.406 | 0.257 | **0.256** | 0.995 | **0.985** | 0.990 | 1.000 | 1.000 |
| $t_1$ | 0.263 | 0.251 | **0.203** | 0.431 | 0.256 | **0.217** | 0.995 | **0.985** | **0.985** | 1.000 | 1.000 |
| $r_a$ | **-0.0009** | 0.008 | 0.008 | **0.015** | 0.017 | 0.018 | 0.960 | 0.960 | **0.950** | 1.000 | 1.000 |
| $\log(t_{12})$ | 0.137 | 0.056 | **0.032** | 0.194 | 0.059 | **0.038** | 1.000 | **0.995** | **0.995** | 1.000 | 1.000 |
| $d_3$ | **0.009** | -0.125 | -0.099 | 0.104 | 0.127 | **0.102** | 0.990 | **0.985** | 0.990 | 1.000 | 1.000 |
| $d_4$ | **0.034** | 0.045 | 0.047 | 0.081 | **0.057** | 0.061 | 0.985 | **0.975** | 0.980 | 1.000 | 1.000 |
| $\log(1 + t_{23})$ | **0.063** | 0.114 | 0.122 | **0.100** | 0.114 | 0.122 | 0.985 | **0.980** | 0.990 | 1.000 | 1.000 |
| $d_{34}$ | **-0.063** | -0.097 | -0.089 | 0.119 | 0.103 | **0.096** | 0.995 | 0.995 | **0.990** | 1.000 | 1.000 |
| $\log(1 + t_{34})$ | 0.086 | **-0.002** | 0.036 | 0.109 | **0.016** | 0.039 | 1.000 | **0.995** | **0.995** | 1.000 | 1.000 |
| $\log(N_a)$ | 0.148 | 0.041 | **0.031** | 0.196 | 0.045 | **0.039** | 1.000 | **0.990** | **0.990** | 1.000 | 1.000 |

